# The ATG5 Interactome Links Clathrin Vesicular Trafficking With The ATG8 Lipidation Machinery For Autophagosome Assembly

**DOI:** 10.1101/769059

**Authors:** Kiren Baines, Jon D. Lane

## Abstract

Autophagosome formation involves the sequential actions of conserved ATG family proteins that regulate the lipidation of the ubiquitin-like modifier ATG8 at the nascent isolation membrane. Although the molecular steps driving this process are well understood, the source of membranes supplied for the expanding autophagosome and their mode of delivery remain uncertain. Here, we have used quantitative SILAC-based proteomics to identify proteins that associate with the ATG12∼ATG5 conjugate that is crucial for ATG8 lipidation. Our datasets reveal a strong enrichment of regulators of clathrin-mediated vesicular trafficking, including clathrin heavy and light chains, and several clathrin adaptors. Also identified were PIK3C2A (a phosphoinositide 3-kinase involved in clathrin-mediated endocytosis) and HIP1R (a component of clathrin vesicles), and the absence of either of these proteins caused defects in autophagic flux in cell-based starvation assays. To determine whether the ATG12∼ATG5 conjugate reciprocally influences trafficking within the endocytic compartment, we captured the cell surface proteomes of autophagy-competent and autophagy-incompetent mouse embryonic fibroblasts under fed and starved conditions. Proteins whose surface expression increased contingent on autophagic capability included EPHB2, SLC12A4, and JAG1. Those whose surface expression was decreased included CASK, SLC27A4 and LAMP1. These data provide evidence for direct regulatory coupling between the ATG12∼ATG5 conjugate and the clathrin membrane trafficking system, and suggest candidate membrane proteins whose trafficking within the cell may be modulated by the autophagy machinery.

## INTRODUCTION

The catabolic macroautophagy pathway (“autophagy”, henceforth) describes the formation de novo of double-membrane-bound vesicles (autophagosomes) that sequester cargo for delivery to the lysosome for degradation. Autophagy occurs at low basal rates in healthy cells but is rapidly upregulated during nutrient/environmental stress. Coordinating the autophagosome assembly process is a family of autophagy related (ATG) proteins that work in concert to effect the lipidation of ubiquitin-like ATG8 proteins (LC3 and GABARAP family members) at the nascent autophagic isolation membrane (e.g. [1]). ATG8 lipidation is controlled by two parallel conjugation systems. In the first, the ubiquitin-like protein ATG12 is constitutively conjugated to the structural protein ATG5, a step that requires ATG7 as the E1-like activating enzyme and ATG10 as the E2-like conjugating enzyme. ATG5 then binds to ATG16L1, forming the ATG12∼ATG5-ATG16L1 complex, which acts as an E3-like lipid conjugation effector complex that recruits and stimulates ATG3, the E2-like conjugating enzyme in the second system. ATG3 regulates the transfer of PE to ATG8, with ATG7 again acting upstream as the E1-like activating enzyme in this process [2]. Importantly, it is anticipated that lipids from diverse sources within the cell become subsumed by or transferred into the expanding isolation membrane, to meet the acute demand for membrane supply/turnover during nutrient stress [3, 4]. Candidate donor sources include: the ER [5]; ER exit sites/ERGIC [6]; the plasma membrane [7]; mitochondria [8]; COPII vesicles [9]; and the Rab11-positive recycling endosome [10]. Identifying the regulatory pathways that coordinate membrane delivery from these diverse sites remains a major challenge for the field.

Analysis of protein interaction networks within the autophagy pathway can indicate novel relationships between autophagosome intermediates and other organelles and/or membrane systems within the cell. Using multiple autophagy proteins as baits, Behrends et al. carried out a comprehensive quantitative appraisal of the autophagy interactome, revealing multiple novel network nodes linking autophagy with other key cellular pathways [11]. In particular, vesicular trafficking factors, including several RABs and RabGAPs, components of the TRAPPIII (trafficking protein particle III) complex, and key constituents of the clathrin complex were identified as candidate interactors of core autophagy molecules [11]. Subsequently, studies from many other groups have highlighted roles for RabGAPs in autophagy [12], with examples including: TBC1D2A [13]; TBC1D14 [14, 15]; TBC1D25 (OATL1) [16]; and TBC1D5 [17]. Similarly, components of the clathrin machinery have been shown to influence autophagy via the establishment of specific endocytic membrane pools incorporating the sole membrane-spanning core autophagy protein, ATG9, and/or ATG16L1 (e.g. [7, 17, 18]). Crucially, clathrin heavy chain (CLTC) has been reported to interact directly with ATG16L1 to coordinate its internalisation into the endocytic compartment [7], while clathrin adaptors AP2A1 and AP2M1 have been shown to interact with ATG9 [17]. These findings complement the reported associations between clathrin complex components and ATG8 proteins in proteomics datasets [11]. Linking these observations, TBC1D5 has been shown to act as a key determinant of ATG9 trafficking during autophagy, regulating the sorting of ATG9 away from the late endosome via differential associations with the retromer component VPS29 and the AP2 clathrin adapter complex [17]. TBC1D14 overexpression impairs autophagy through the generation of enlarged, tubulated recycling endosomes that retain the autophagy initiation kinase, ULK1 [14], and its influence on autophagy appears to be via binding to TRAPPIII—identified as an interactor of TECPR1 by Behrends et al. [11]. Together, these proteins regulate RAB1-mediated release of ATG9 from the RAB11-positive compartment via the Golgi in an ULK1-independent manner to establish the ATG9-positive vesicular pool [15]. The trafficking of ATG9A from the recycling endosome into pre-autophagic structures that are also positive for ATG16L1 and WIPI2 depends on the sorting nexin SNX18 and dynamin-2 (DNM2) [19, 20], and recently, a role for ARFIP2 in mobilisation of ATG9A-positive vesicles that deliver PI4KIIIbeta to the site of autophagosome assembly has been described [21]. Notably RAB11A and transferrin receptor-positive compartment appear to be pivotal for autophagosome assembly, as here RAB11A binds WIPI2 to establish a platform for autophagosome assembly [10]. Clearly, the autophagy and intracellular membrane trafficking pathways are intimately linked, with routing of ATG9 through the endocytic system a prominent regulatory nexus [22, 23].

While there are numerous examples of how altered intracellular membrane trafficking pathways can impact on the autophagy response, the idea that certain autophagy proteins directly influence the specificity and/or efficiency of protein trafficking through the biosynthetic and/or endocytic routes has gained traction [24]. Autophagy proteins are implicated in unconventional protein secretion [25, 26] with cargoes including the inflammatory mediator IL-1β [27] and the cystic fibrosis ΔF508 CFTR protein [28]. The autophagy machinery also influences the steady state distribution of a subset of surface proteins including the FAS receptor [29] and the glucose transporter, SLC2A1 (GLUT1) [30]. Notably, surface levels of SLC2A1 were elevated in an autophagy-dependent manner in response to increased glucose demand through autophagy-mediated sequestration of TBC1D5 and subsequent diversion of SLC2A1 to the plasma membrane [30]. The autophagy machinery—exemplified by ATG5—also influences the properties of major histocompatibility complex (MHC) class II loading compartments [31, 32], and elevated surface levels of MHC class I have been reported in ATG5 and ATG7 deficient dendritic cells, caused by an internalisation defect linked to differential mobility of AP2-associated kinase (AAK1) [33]. From such studies, the concept of secretory autophagy has emerged [24], and indeed one report using ATG5^flox/flox^ mouse macrophages has provided evidence that a number of secreted, leaderless proteins (including FTH1 [ferritin heavy chain-1] and several lysosomal proteins) have altered, relative extracellular abundances upon ATG5 ablation [34]. Based on these studies, a direct role for autophagosomes in the sequestration and delivery of secreted cargoes has been proposed [34]. With respect to endocytosis, and in addition to the aforementioned links between core autophagy proteins and the clathrin machinery, a novel pathway for association of ATG8 proteins with endosomes has been described in the context of Aβ uptake in a mouse Alzheimer’s disease model [35]. This pathway of LC3-associated endocytosis (LANDO) requires ATG5 but not FIP200 (necessary for canonical autophagy), and curiously is also dependent on Rubicon—a protein that restricts canonical autophagy [35].

Here, we have carried out unbiased quantitative proteomics analyses to characterise the ATG5 interactome in autophagy-competent and -incompetent cells, providing clear evidence for interactions between the lipidation machinery and components of clathrin-dependent membrane trafficking pathways. Indeed, analysis of clathrin adaptors present in our networks suggest that the AP2 and AP1 clathrin systems converge on ATG5. Within the clathrin endocytic machinery identified in the ATG5 interactome, we provide evidence suggesting that two putative ATG5 interactors—PIK3C2A and HIP1R—influence autophagic flux in vitro. Finally, using surface biotinylation, we capture the surface proteomes of autophagy-competent and -incompetent cells under full media and starvation conditions, as a step towards understanding how the autophagy machinery influences conventional trafficking through the endocytic and secretory systems.

## RESULTS & DISCUSSION

### Quantitative ATG5 proteomics describes the hierarchical recruitment of autophagy regulators at the isolation membrane

To identify novel proteins that are recruited to the isolation membrane at the point of membrane expansion and shaping, we used GFP-TRAP immunoisolation coupled with SILAC (stable isotope labelling with amino acids in culture)-based quantitative proteomics to define the ATG5 interactome. We used lentiviruses to stably “rescue” ATG5^−/−^ MEFs with: (i) GFP; (ii) wild-type GFP-ATG5 (WT GFP-ATG5); or (iii) K130R GFP-ATG5 (a point mutation that prevents conjugation to ATG12 [36]). These constructs were chosen to enable comparative analyses of the interactomes of conjugation competent and conjugation deficient ATG5. In ATG5^−/−^ MEFs, MAP1LC3B (LC3B) lipidation was absent as expected, and the levels of SQSTM1/p62 (a protein that is degraded by autophagy), were constitutively high (**Supplemental Fig. S1A**). Consistent with this, LC3B puncta were dramatically reduced (even when cells were starved in the presence of Bafilomycin A1 [BafA1]), while WIPI2 puncta numbers were elevated even under basal conditions (**Supplemental Fig. S1B-D**). Rescued MEFs were validated for their responses to amino acid/growth factor starvation (1 hour), and as anticipated, only the WT GFP-ATG5 cell-line supported LC3B lipidation (**Fig. 1A**), showed normal WIPI2 puncta responses (**Fig. 1B**), and assembled LC3B-decorated autophagosomes (**Fig. 1C**). In both the WT GFP-ATG5 and K130R GFP-ATG5 cell-lines, WIPI2 co-localised with GFP-ATG5 at presumed autophagosome assembly sites during starvation (**Fig. 1D**). In K130R-GFP-ATG5 cells, these structures were confirmed as stalled isolation membranes— flattened, electron-dense membrane structures cradled by characteristic ER tubules [37, 38]— using correlative light and electron microscopy (CLEM) (**Fig. 1E**).

**Fig. 1.**
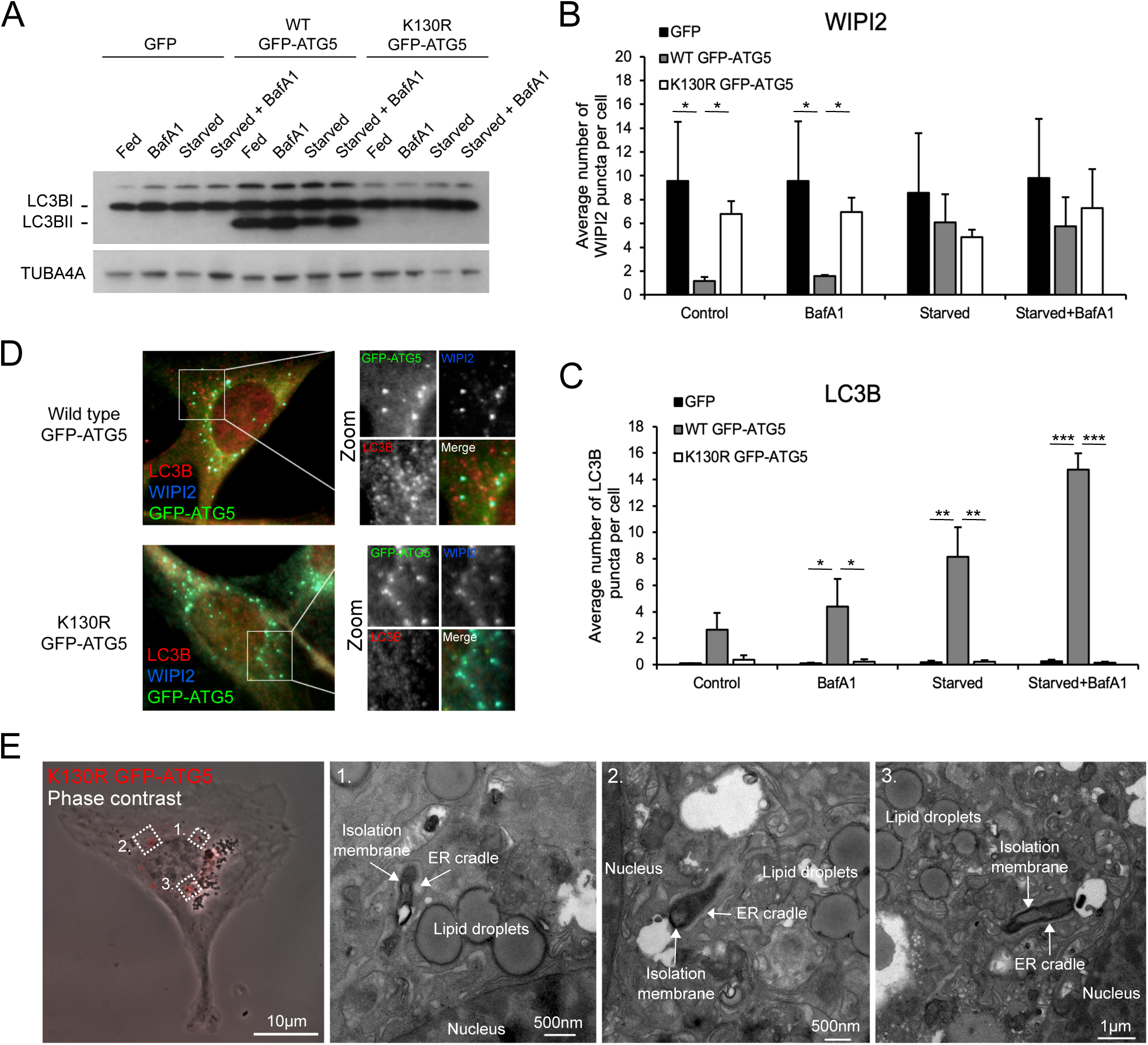
Characterisation of cell-lines used for SILAC proteomics. (A) Immunoblot of ATG5^−/−^ MEFs stably expressing GFP, wild-type (WT) GFP-ATG5 or K130R GFP-ATG5. Cells were starved for 1 hour, and lysates probed for LC3B and TUBA4A. (B) WIPI2 puncta counts in fed and starved cells, in the absence or presence of BafA1. (C) LC3B puncta counts in fed and starved cells, in the absence or presence of BafA1. (D) Example images of ATG5^−/−^ MEFs stably expressing WT GFP-ATG5 or K130R GFP-ATG5, fixed and stained with antibodies against WIP2 and LC3B. Bar = 10 *µ*m. (E) CLEM analysis of K130R GFP-ATG5 MEFs. Mean ± SEM; n = 3; 10 cells per condition per experiment; statistical significance calculated using ANOVA, followed by Tukey’s HSD test. (***P < 0.001, **P < 0.01, *P < 0.05).

Each MEF cell-line was then grown in SILAC media for 8 cell doubling events, before being starved for 1 hour, and then lysed for GFP-Trap immunoisolation (**Fig. 2A**). Samples were analysed by mass spectroscopy, and a Sequest search carried out against the Uniprot mouse database with a <5% FDR cut-off applied to eliminate low confidence hits (see **Materials & Methods**). Positive interactors were identified as those that were present in the WT GFP-ATG5 interactome but were absent in the GFP equivalent (2 or more individual peptides; >2-fold enrichment over GFP), and the obtained dataset comprised of 495 proteins (**Supplemental Table 1** showing representative data from 2 identical experiments). Of these, 218 proteins were also enriched >2-fold over the K130R GFP-ATG5 dataset, and these represented candidate ATG12∼ATG5 conjugate interactors (**Supplemental Table 2**). Here, there was a clear enrichment of proteins involved in autophagosome assembly, as expected. Similar analysis of the K130R GFP-ATG5 dataset revealed 364 proteins enriched >2-fold over GFP, of which 127 proteins also showed a >2-fold enrichment over WT GFP-ATG5, and these represented candidate unconjugated ATG5 interactors (**Supplemental Table 3**).

**Fig. 2.**
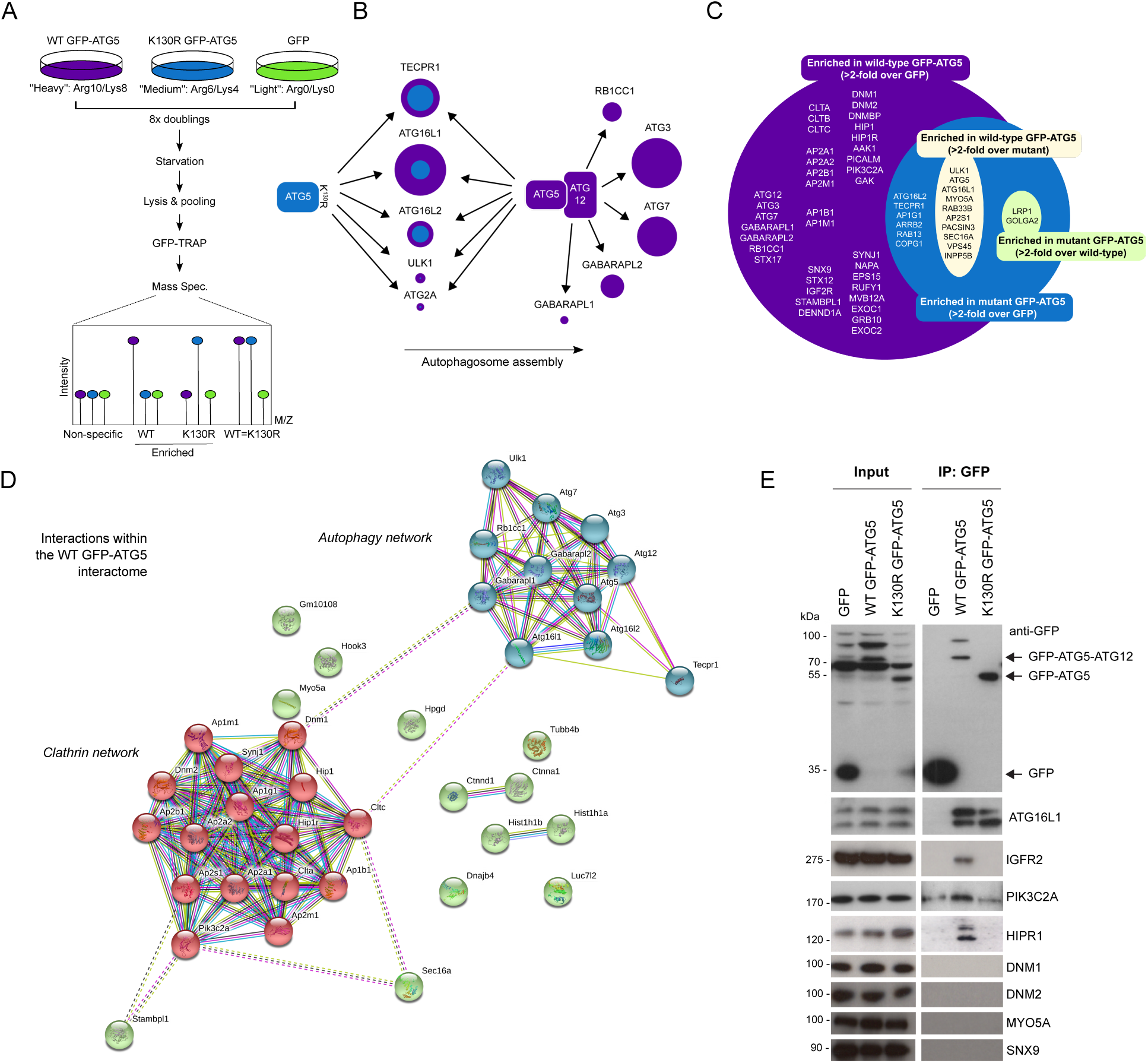
Analysis of the ATG5 interactome. (A) SILAC protocol schematic. (B) Cartoon showing the autophagy protein interactions identified in the WT GFP-ATG5 and K130R GFP-ATG5 interactomes. Circles representing proteins are sized by their SILAC “score” as an indicator of confidence within the dataset. Proteins common to both datasets are represented by magenta (WT GFP-ATG5) and blue (K130R GFP-ATG5) circles. The relative areas of the two provide a rapid visual depiction of SILAC relative abundance ratios. (C) Venn diagram mapping the interactions within the different groups that are classified as having roles in autophagy or membrane trafficking. (D) STRING analysis of the high confidence WT GFP-ATG5 interactome showing 2 distinct interaction hubs: autophagosome biogenesis and clathrin-mediated vesicular trafficking. (E) Immunoblots of selected candidate interactors. GFP-TRAP immunoisolates from the GFP, WT GFP-ATG5 and K130R GFP-ATG5 MEFs under starvation conditions (1 hour).

Analysis of proteins identified in the WT GFP-ATG5 and K130R GFP-ATG5 datasets pointed to a hierarchical recruitment of autophagosome assembly factors in autophagy competent and incompetent cells, which broadly agreed with expected recruitment kinetics at autophagosome assembly sites [39, 40] (**Fig. 2B**). For example, proteins acting during early stages of autophagosome assembly were detected >2-fold over GFP in the unconjugated K130R GFP-ATG5 data (see **Supplemental Table 2**; e.g. ATG16L1; ATG2A; ULK1); whereas proteins that were enriched in the WT GFP-ATG5 interactome (**Supplemental Table 2**; i.e. >2-fold over K130R GFP-ATG5) included those linked to the latter stages of autophagosome maturation and ATG8 lipidation (e.g. ATG3; ATG7; GABARAP-L1; GABARAP-L2/GATE-16) (**Fig. 2B**). Several other ATG8 family members were detected in the SILAC datasets, but below the peptide number threshold (namely: GABARAP; LC3A; LC3B; data not shown).

Unexpectedly, TECPR1, a protein that has been shown to bind directly to the ATG12∼ATG5 conjugate to facilitate autophagosome-to-lysosome fusion [41], was found to be a strong interactor in both WT GFP-ATG5 and K130R GFP-ATG5 over GFP datasets (**Supplemental Table 1**). This might indicate additional roles for TECPR1 during early stages of autophagosome formation, and indeed, TECPR1 has been observed to colocalise with ATG5 at presumed autophagosome assembly sites in co-overexpressing cells [41]. It was also interesting that ATG16L1 was enriched in both the wild type and K130R GFP-ATG5 datasets (**Fig. 2B**; **Supplementary Table 2**), as this suggested an association with unconjugated ATG5. Indeed, ATG16 has been found to bind directly to [42] and reinforce the lipid binding capabilities [43] of ATG5, although ATG16L1 is dispensable for the E3-like activity of the ATG12∼ATG5 conjugate in vitro [2]. Of further interest, K130R GFP-ATG5 also interacted strongly with ATG16L2—an isoform of mammalian ATG16 that binds to ATG12∼ATG5 but cannot support autophagosome assembly due to variations in the coiled coil central section [44]. Interestingly, RBCC1 (also known as FIP200) was only identified in the WT GFP-ATG5 dataset [**Fig. 2B**; **Supplementary Table 2**]). Other components of the mammalian ULK1 complex (namely, ATG13 and ATG101) were not found in any of the datasets. Additional autophagosome assembly factors that were detected below the 2-fold enrichment threshold over GFP in the WT GFP-ATG5 dataset included: ATG2B; ATG10; SQSTM1/p62; FYCO1; WIPI3; WIPI4 (data not shown).

### Enrichment of the clathrin trafficking machinery in the WT GFP-ATG5 interactome

In addition to the core autophagy-associated proteins identified in the two GFP-ATG5 interactomes (**Supplementary Tables 2, 3**), there was a substantial enrichment of proteins with roles in membrane trafficking (**Fig. 2C**). Of those enriched in the WT GFP-ATG5 interactome, a high proportion identified as being involved in endocytic trafficking, and in particular clathrin-mediated vesicle trafficking (**Supplementary Table 2; Fig. 2C**). The majority of these proteins were absent in the K130R GFP-ATG5 dataset (**Supplementary Table 3**), suggesting that ATG12∼ATG5 conjugation is needed for this interaction (examples included: clathrin heavy chain [CLTC]; clathrin light chain [CLTA]; sorting nexin 9 [SNX9]; huntingtin interacting protein 1 [HIP1]; huntingtin interacting protein 1 related [HIP1R]; cyclin G-associated kinase [GAK]; dynamin binding protein [DNMBP]; AP2-asssociated kinase 1 [AAK1]; epidermal growth factor receptor substrate 15 [EPS15]; and phosphatidylinositol 4-phosphate 3-kinase C2 domain-containing subunit alpha [PIK3C2A]). High confidence interactome lists were next assembled to include only those proteins that were >8-fold enriched (vs. GFP), with >7.5 Score values. These were subjected to STRING analysis to visualise relationships within protein families (**Fig. 2D**). Evident from this analysis for the WT GFP-ATG5 dataset were two clearly separate interaction groups: (i) core autophagosome assembly proteins; (ii) regulators of clathrin-dependent vesicular trafficking (**Fig. 2D**). By contrast, aside from the relatively small number of autophagy proteins that were present, the only other possible protein grouping that emerged from parallel K130R GFP-ATG5 STRING analysis included components of the extracellular matrix (**Supplemental Fig. S2**). In addition to clathrin heavy and light chains reported in the WT GFP-ATG5 dataset, a number of clathrin adaptor subunits of the AP1 (B1; M1; G1) and AP2 (A1; A2; B1; M1) classes were identified (**Fig. 2D**). Overall, the identification of proteins involved in the control of membrane phosphoinositide identity (PIK3C2A; SYNJ1), mediators of clathrin recruitment/activity during coat assembly/scaffolding (PIK3C2A; SNX9; PICALM; AAK1; EPS15) and uncoating (GAK), clathrin adaptor and local actin regulator (HIP1R), and regulators of membrane deformation/scission (DNM1; DNM2; SNX9) at the site of endocytic uptake, suggests that ATG5 interacts with the clathrin-associated endocytic machinery [45]. The presence of AP1 adaptor components also points to a possible relationship between ATG5 and the Golgi-to-early/recycling endosome trafficking step [46], meanwhile the enrichment of insulin-like growth factor 2 receptor (IGFR2; also known as the cation-independent mannose-6-phosphate receptor, CI-M6PR), which delivers hydrolytic enzymes to the lysosome, argues for a more global role in clathrin-mediated trafficking within the cell.

The identification of CLTC in our data and in the Behrends et al. ATG5 interactome [11] was of particular interest since previous research has shown that ATG16L1 can be co-isolated with CLTC (and AP2), and that CLTC contributes to autophagosome biogenesis via the establishment of a distinct ATG16L1-positive endocytic membrane compartment [7, 18]. How ATG5—either alone or in its ATG12-conjugated form—contributes to this process remains uncertain. Experimental data suggest that the N-terminal domain of ATG16L1 is responsible for the CLTC interaction, but given that the ATG12∼ATG5 conjugate also binds to ATG16L1 in this region, it is uncertain how ATG12∼ATG5 influences this association (although overexpression of ATG5 was found not to disturb the ATG16L1/CLTC interaction) [7]. The presence of ATG16L1 in both our WT GFP-ATG5 and K130R GFP-ATG5 datasets, suggests that binding is not contingent on ATG12 conjugation (**Supplementary Table 2; Fig. 2B**). Since CLTC, and in fact the large majority of the other clathrin-mediated endocytosis-associated proteins identified in our datasets, were absent in the K130R GFP-ATG5 interactome (**Supplementary Table 3; Fig 2C**), this suggests it is unlikely that ATG5 engagement with the clathrin machinery occurs via ATG16L1. It would be interesting to test whether the novel surface patch is established when the ATG12∼ATG5 conjugation is formed (see s[42]), facilitates interactions with proteins such as those associated with the clathrin machinery.

### Validation of candidate ATG12∼ATG5 interactors

We next sought to validate selected candidate interactors enriched in the WT GFP-ATG5 interactome, namely: PIK3C2A; HIP1R; IGFR2; SNX9; MYO5A; DNM1; DNM2 (**Fig. 2C**). GFP-trap pull-downs of lysates from the GFP, WT GFP-ATG5, and K130R GFP-ATG5-expressing ATG5^−/−^ MEFs were analysed by immunoblotting. This clearly showed the presence of ATG16L1 in both WT GFP-ATG5 and K130R GFP-ATG5 pull-downs (**Fig. 2E**), as predicted from the SILAC-based proteomics analysis (**Supplementary Table 2**). By contrast, PIK3C2A, HIP1R and IGFR2 were strongly enriched in the WT GFP-ATG5 pull-down lysate only (**Fig. 2E**). However, MYO5A, DNM1, DNM2, and SNX9 could not be detected in immunoblots of pull-downs from either WT or K130R GFP-ATG5 cells (**Fig. 2E**).

To establish whether the link between ATG12∼ATG5 and the clathrin system is dependent upon productive autophagosome assembly, we established cell-lines stably expressing GFP, WT GFP-ATG5 or K130R GFP-ATG5 in an ATG3^−/−^ MEF background [47] for SILAC-based proteomics (**Supplementary Fig. S3**). As expected, ATG3^−/−^ MEFs were deficient in LC3B lipidation and autophagy, even when supplemented with the ATG5 constructs (**Supplementary Fig. S3A, B**). Quantitative proteomics was carried out following nutrient starvation as before; proteins with a peptide count <2 were removed, and a WT GFP-ATG5 interactome was constructed comprising proteins that were >2-fold enriched over GFP and K130R GFP-ATG5 (**Supplementary Tables 4, 5**). Overall, the comparable level of enrichment for WT GFP-ATG5 interactors identified in the ATG3^−/−^ background was lower than in the ATG5^−/−^ background, perhaps because GFP-ATG5 was expressed alongside the endogenous protein. Core autophagy proteins that were enriched in the WT GFP-ATG5 interactome in ATG3 null MEFs included: GABARAPL2 (GATE-16); ATG7; ATG12; TECPR1; ATG16L1; and ATG16L2 (ATG3 peptides detected are likely to be the truncated product of the mutant ATG3 allele generated by targeting exon 10 [47]). Also identified were several proteins implicated in the regulation of clathrin-mediated trafficking, including: CLTC; CLTA; HIP1R; SNX9; DNM1; DNM2; PIK3C2A (**Supplementary Table 4**). STRING analysis of this dataset once again revealed 2 clusters centred on autophagy and clathrin-mediated endocytic trafficking (**Supplementary Fig. S3C**). Similar analysis of the K130R GFP-ATG5 interactome once again only showed evidence of a limited cluster of autophagy proteins (**Supplementary Fig. S3D**). In this background, immunoblotting confirmed that ATG16L1 co-isolated with both WT GFP-ATG5 and K130R GFP-ATG5, whereas PIK3C2A was only detected in WT GFP-ATG5 pull-downs (**Supplementary Fig. S3E**). The co-isolation of these proteins in the ATG3^−/−^ background argues that ATG12∼ATG5 can still engage with clathrin-associated trafficking factors regardless of the requirement for progressive autophagosome assembly and the accompanying need for membrane supply at autophagosome assembly sites.

### PIK3C2A and HIP1R contribute to basal and starvation-induced autophagy

To determine the possible relevance of interactions between the ATG12∼ATG5 conjugate and proteins linked to clathrin-mediated endocytosis, we analysed the autophagy responses in cells lacking either PIK3C2A or HIP1R. Phosphatidylinositol 3-kinases (PI3Ks) are lipid kinases involved in a large set of biological processes, including endocytic trafficking, receptor signalling, organization of the cytoskeleton, and autophagy [48, 49]. There are three PI3K classes and 8 PI3K proteins in total, all of which phosphorylate the D3 position of the inositol ring of phosphatidylinositol (PIs) to generate PI3P [50], with the sole class III member, VPS34, being responsible for a major fraction of PI3P within cells needed during autophagosome formation [51]. Of the less well characterised class II PI3Ks (PIK3C2A; PIK3C2B; PIK3C2C), PIK3C2A can also form phosphatidylinositol-3,4-bisphosphate (PI(3,4)P_2_ [52], a lipid that is concentrated at the plasma membrane and is required for the nucleation of clathrin-coated pits (e.g. [53]). At the plasma membrane, PIK3C2A spatiotemporally controls clathrin-mediated endocytosis by regulating clathrin-coated pit maturation [52]. PIK3C2A plays a crucial role in development as loss of the *Pik3c2a* gene in mouse causes early embryonic lethality due to defective vascular development [54] and homozygous loss-of-function mutations have recently been identified in patients with broad-ranging developmental abnormalities linked to ciliary defects [55]. Indeed, PIK3C2A influences cilia formation by regulating sonic hedgehog signalling [56]. In addition to these roles, data suggest that PIK3C2A may have a role in autophagy: it was identified as an interactor of ATG7 (but not ATG5) in the Behrends et al. study [11], and a reduction in LC3 puncta numbers has been observed when PIK3C2A is silenced together with PIK3C2B [48]. This prompted us to further explore the involvement of PIK3C2A in autophagy control.

PIK3C2A^flox/flox^ MEFs (kindly provided by Prof. Yoh Takuwa; Kanazawa University [54]) were used to assess the formation of autophagosomes in basal and starvation conditions (**Fig. 3**). Immunofluorescence staining for WIPI2 and LC3B was carried out with or without addition of Cre recombinase (48 hours), which caused efficient ablation of PIK3C2A expression (**Fig. 3A, B**). Quantitation revealed that removal of PIK3C2A led to a significant increase in LC3B puncta numbers in full nutrient conditions, and a further increase upon addition of BafA1, consistent with elevated basal autophagic flux (**Fig. 3A, D**). Accordingly, WIPI2 puncta numbers were also elevated in Cre-treated cells in basal conditions (statistically significant in the BafA1-treated population; **Fig. 3A, C**). Interestingly, this pattern was not repeated in starved cells: here, LC3B and WIPI2 puncta numbers were similar in the absence and presence of Cre (**Fig. 3A, D**). Interestingly, addition of BafA1 to nutrient starved cells caused no further increases in LC3B or WIPI2 puncta numbers, indicative of a block in autophagic flux (**Fig. 3A, D**). This suggests that PIK3C2A may suppress autophagy in basal conditions whilst contributing to the stimulation of a robust autophagic flux response during nutrient starvation.

**Fig. 3.**
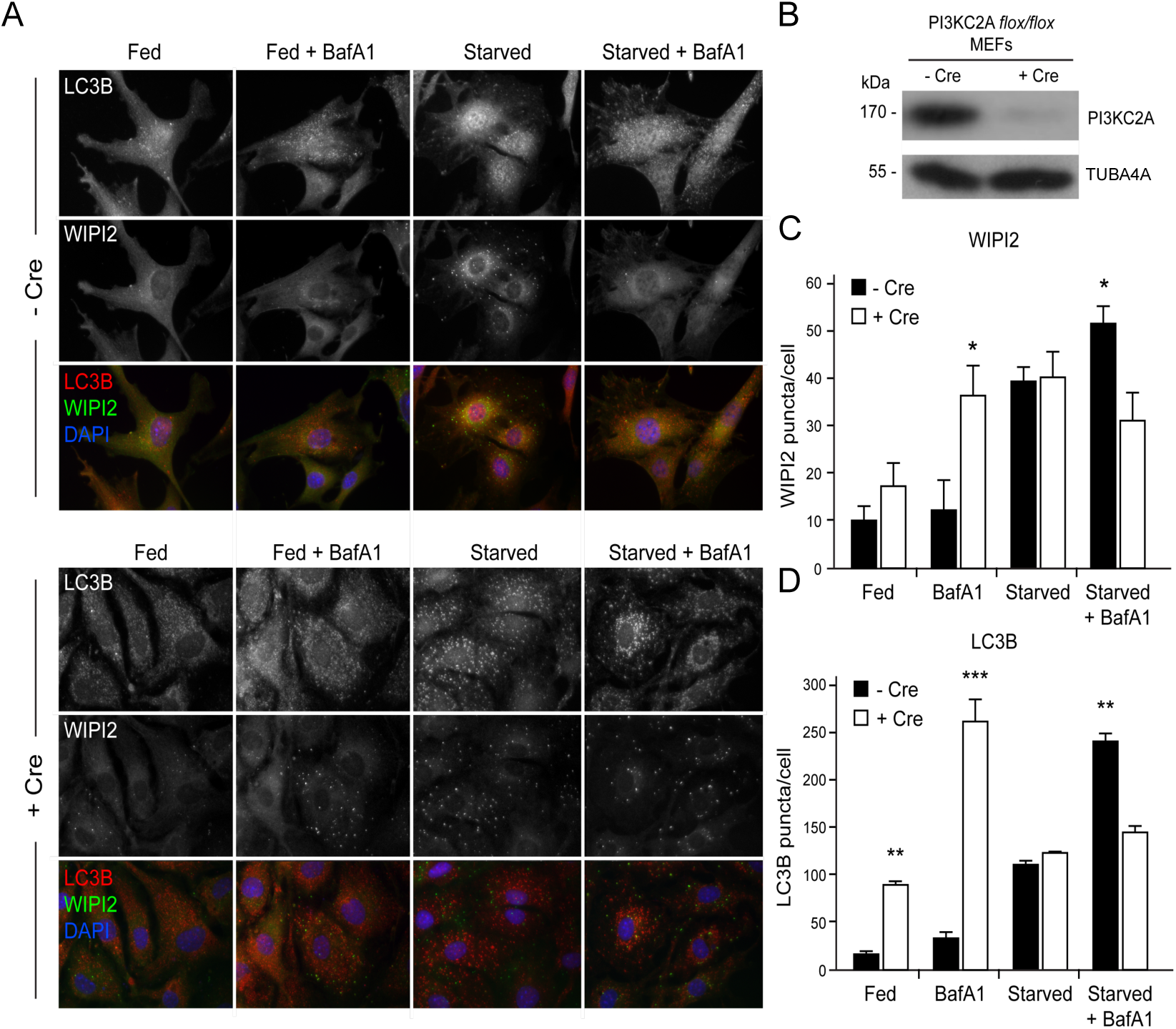
PIK3C2A suppresses basal autophagy but is needed for a robust starvation response. (A) Immunofluorescence images of PIK3C2A^flox/flox^ MEFs under full nutrients or starved in the absence or presence of BafA1, with or without addition of Cre recombinase. Cells were fixed then stained with antibodies against LC3B (red) and WIPI2 (green). DAPI staining is in blue. Bar = 10 *µ*m. (B) Immunoblot of PIK3C2A^flox/flox^ MEFs with or without Cre recombinase. (C) WIPI2 puncta quantitation. (D) LC3B puncta quantitation. Mean ± SEM; n = 3; 10 cells per condition per experiment; statistical significance calculated using ANOVA, followed by Tukey’s HSD test. (***P < 0.001, **P < 0.01, *P < 0.05).

HIP1R is one of a number of essential adaptor proteins that regulate clathrin-mediated endocytosis. It is a component of clathrin-coated pits and binds to both CLTA and actin filaments [57, 58]. HIP1R has also been shown to have a role in vesicular trafficking at the TGN [59]. CLTA plays a crucial role in recruiting HIP1R to clathrin-coated pits, via the central dimerization-competent coiled-coil region of HIP1R, which in turn regulates local actin assembly [60–63]. Furthermore, HIP1R has an N-terminal ANTH (AP180 N-terminal homology) domain, through which it interacts with phospholipids [64, 65]. Since both the clathrin-mediated endocytic system and the actin cytoskeleton contribute to the autophagosome assembly [7, 66], HIP1R may couple these systems to the autophagy machinery via ATG12∼ATG5 (**Fig. 2C-E**). Wild-type and HIP1R null MEFs (kindly provided by Dr. Theodora Ross; UT Southwestern Medical Centre) were used to assess the contribution of HIP1R during basal and starvation-induced autophagy by immunofluorescence staining for WIPI2 and LC3B (**Fig. 4**). The autophagy response of HIP1R null cells was indicative of a strong block in autophagic flux: in basal conditions, LC3B puncta numbers were significantly higher than in wild-type MEFs, but did not increase substantially in the presence of BafA1 (**Fig. 4A, D**). No further increase was recorded following nutrient starvation in the absence of BafA1, however in the presence of BafA1 there were significantly fewer LC3B puncta numbers compared to wild-type cells (**Fig. 4A, D**). WIPI2 puncta numbers in HIP1R null MEFs followed a similar pattern to wild-type cells, although in the starved + BafA1 condition WIPI2 puncta numbers were significantly lower in HIP1R null cells (**Fig. 4A, C**).

**Fig. 4.**
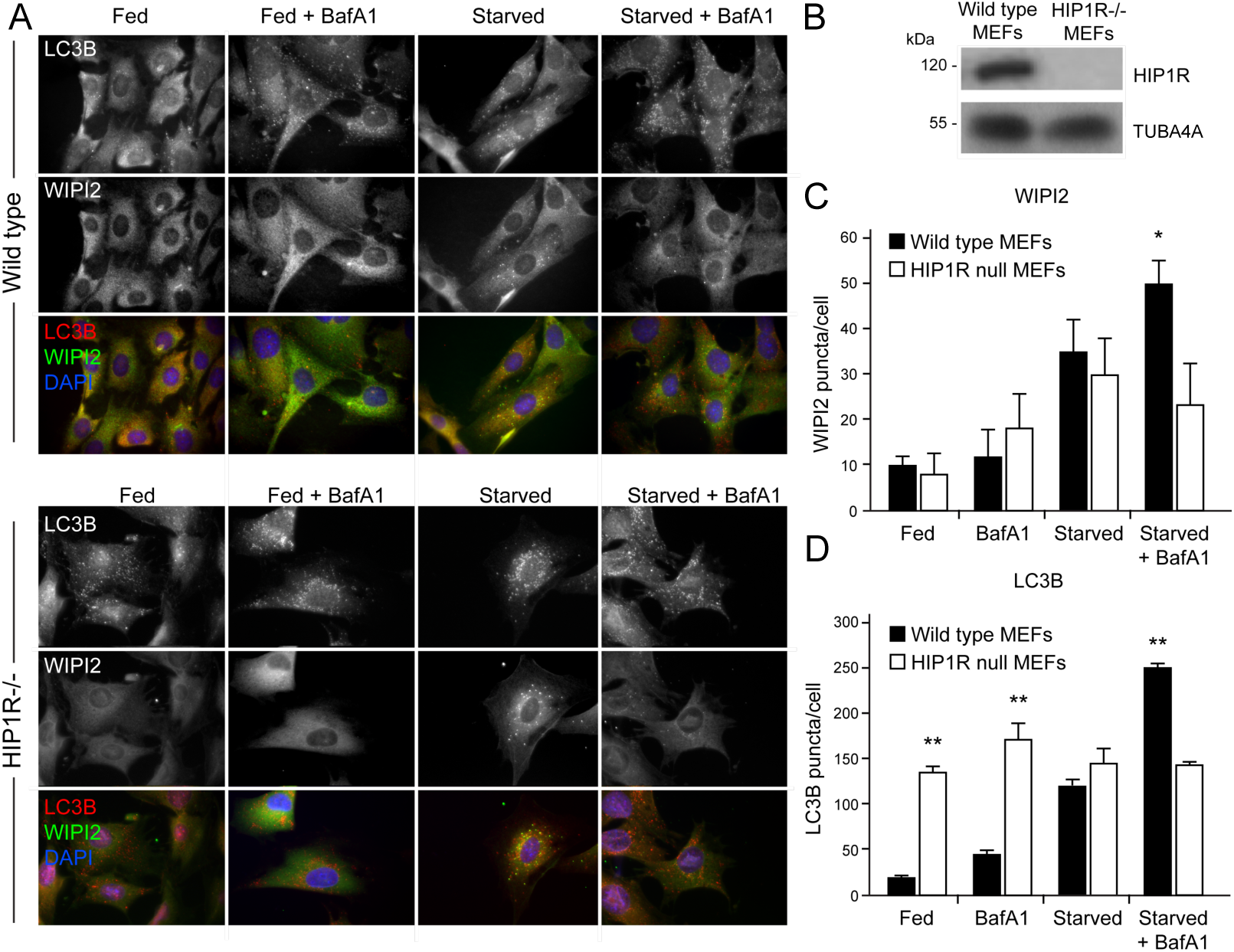
Absence of HIP1R stimulates basal autophagy but suppresses the autophagy response to starvation. (A) Immunofluorescence images of wild-type and HIP1R^−/−^ MEFs under full nutrients or starved in the absence or presence of BafA1. Cells were fixed then stained with antibodies against LC3B (red) and WIPI2 (green). DAPI staining is in blue. Bar = 10 *µ*m. (B) Immunoblot wild-type and HIP1R^−/−^ MEFs. (C) WIPI2 puncta quantitation. (D) LC3B puncta quantitation. Mean ± SEM; n = 3; 10 cells per condition per experiment; statistical significance calculated using ANOVA, followed by Tukey’s HSD test. (***P < 0.001, **P < 0.01, *P < 0.05).

### Quantitative SILAC cell surface proteomics and the ATG12∼ATG5 conjugate

The proteomics data linking ATG12∼ATG5 and the clathrin-mediated vesicle trafficking machinery prompted us to test whether the steady state surface proteome is influenced by the actions of the ATG12∼ATG5 conjugate in supporting a functional autophagy system. We used biotinylation to capture the surface proteomes of the reconstituted ATG5^−/−^ MEFs (GFP; WT GFP-ATG5; K130R GFP-ATG5) under fed and starved conditions (respectively, 4 and 3 independent surface biotinylation experiments). Firstly, to assess the cell surface proteome under fed conditions, cells were grown in SILAC media as before (**Fig. 2A**), subjected to surface biotinylation, lysis and streptavidin pull-down, then processed by SILAC-based quantitative proteomics. Following data sampling to exclude low confidence peptides, and data normalisation to account for variabilities in labelling efficiency (see **Materials & Methods**), proteins that were represented in at least 3 of the 4 repeats were identified and their relative abundancies compared between the three different cell-lines (**Supplemental Tables 6-9**).

Data obtained from cells under full nutrient conditions were displayed as volcano plots (log10 mean vs -log10 p-value; **Fig. 5A-C**). Individual proteins that were consistently altered in the WT GFP-ATG5 vs. GFP dataset included: EPHB2; PVRL1/NECTIN-1; PTPRF-PE; JAG1; HNRNPLL; ATP6V1B2; SLC12A4 (all up); and LXN; STIM1; CASK; SLC27A4 (FATP4) (all down) (**Fig. 5A; Supplemental Tables 6, 7**). JAG1 is involved in NOTCH signalling, while PVRL1 and EPHB2 are surface proteins with roles in cell-to-cell adhesion, cell migration and cell repulsion/adhesion. Interestingly, NOTCH signalling has been shown to link with the autophagy pathway in recent studies [67–69]. Examples of proteins that differed between the WT GFP-ATG5 and K130R GFP-ATG5 datasets included: DCHS1; VLDLR; PLXNA2; VLDLR; SLC12A2 (all up); and STIM1; TPD52L2; PON3; SLC27A4; H2-D1; MAP2K1 (all down) (**Fig. 5B; Supplemental Tables 6, 8**). Once again levels of SLC27A4 were reduced in the WT GFP-ATG5 surface proteome relative to autophagy-deficient cells. SLC27A4 is a member of the fatty acid transport protein family with roles in regulated fatty acid uptake [70], and its relative enrichment on the surface of autophagy-deficient cells might be expected in cells compensating for the lack of autophagy-mediated fatty acid mobilisation. Interestingly, recent data suggests a possible link between SLC27A4 and ATG4B in lung cancer cells [71]. SLC12A2 (also known as NKCC1) is a Na^+^/K^+^/2Cl^−^ transporter that regulates cellular ion levels and volume, linking cell volume control and mTORC1 signalling [72]. Comparing the K130R GFP-ATG5 and GFP surface proteomes revealed differences in levels of: ICOSLG; ABI3BP; DECR1; MLYCD; NT5DC1; SSR4 (all up); and SLC16A3 (MCT4); LAMB1; CLMP; PTK7 (all down) (**Fig. 5C; Supplemental Tables 6, 9**). SLC16A3 is a monocarboxylate transporter that plays key roles in metabolism through the uptake of substrates including pyruvate and L-lactate, as well as ketone bodies, a range of α-keto monocarboxylates, and short chain fatty acids [71, 73].

**Fig. 5.**
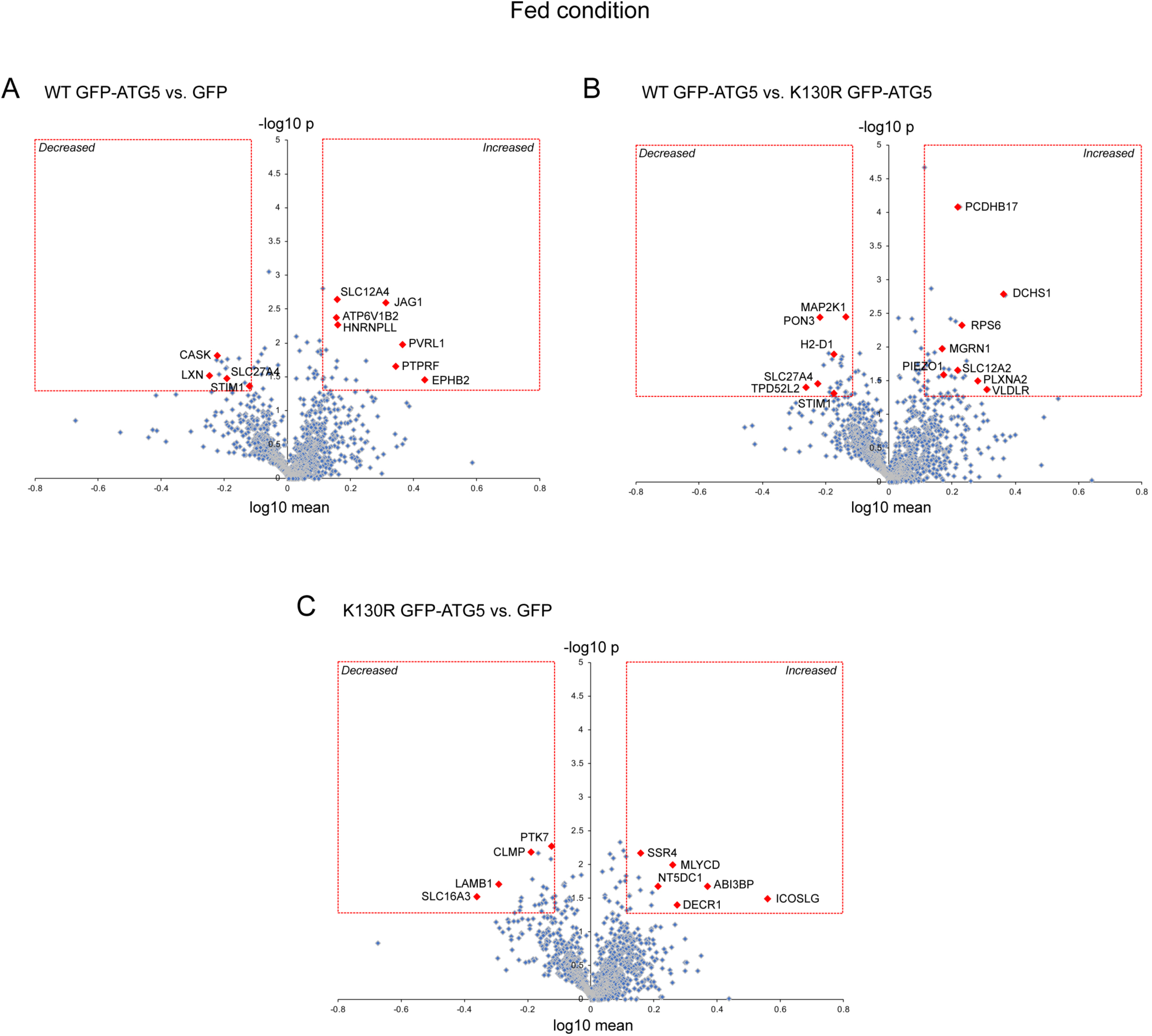
Changes in basal surface protein abundance in ATG5^−/−^ MEF rescued with GFP, WT GFP-ATG5 or K130R GFP-ATG5. Pairwise comparisons of surface protein abundance analysed by surface biotinylation/streptavidin pull-down and SILAC quantitative proteomics in fed conditions. (A) WT GFP-ATG5 vs. GFP. (B) WT GFP-ATG5 vs. K130R GFP-ATG5. (C) K130R GFP-ATG5 vs. GFP. In all examples, the red boxes indicate 1.3-fold enrichment cut-off and p<0.05. Red diamonds depict selected proteins of potential interest.

We next analysed the surface proteomes of cells subjected to 1h nutrient starvation (**Fig. 6A-C; Supplemental Tables 10 & 11**). From the 3 proteomics datasets obtained, 494 (WT GFP-ATG5 vs. GFP), 609 (WT GFP-ATG5 vs. K130R GFP-ATG5) and 409 (K130R GFP-ATG5 vs. GFP) proteins were represented on each occasion (**Supplemental Table 10**). Of those, proteins whose levels were significantly changed >1.3-fold on the surface of WT GFP-ATG5 over GFP-expressing cells included: ROBO1; PARK7; TSPAN15; MTHFD1L (all up); and COL12A1; BST2; SYPL; LAMP1, SLC6A6, STX12; APP (all down) (**Fig. 6A; Supplemental Table 11**). Comparing WT GFP-ATG5 against K130R GFP-ATG5 surface proteomes revealed that no proteins were significantly increased, whereas those whose levels were reduced >1.3-fold included: LAMP1; BST2; SLC27A4; APP; LSS; LMAN1; LRPAP1 (**Fig. 6B; Supplemental Table 11**). The absence of proteins increasing in this pairwise comparison, and the duplication of certain proteins decreasing when comparing WT GFP-ATG5 with both GFP and K130R GFP-ATG5 (namely, LAMP1; BST2; APP) (**Fig. 6A, B; Supplemental Table 11**), is indicative of a general reduction of surface protein levels during starvation in autophagy-competent cells. Comparing K130R GFP-ATG5 against GFP revealed the following proteins that significantly changed following starvation: LAMP1; HK1; MTHFD1L; SACM1L; STT3A (all up); and MOB1B; CAMK2D; ERBB2; ACACA (all down) (**Fig. 6C; Supplemental Table 11**). Of these proteins, LAMP1 stood out as being significantly decreased in the WT GFP-ATG5 surface proteomes when compared with both GFP and K130R GFP-ATG5 datasets. One interpretation of this is that LAMP1—an important lysosomal glycoprotein—is preferentially internalised during starvation in autophagy-competent cells, and that the presence of conjugation-deficient ATG5 impairs this process (**Fig. 6A-C; Supplemental Table 11**). Similarly, reduced surface expression of BST2 (also known as tetherin/CD317—a viral restriction factor and exosome tether that associates with lipid rafts via tandem *trans*-membrane domain and GPI anchor [74–76]) might suggest a link between autophagy competency and viral and/or exosome release.

**Fig. 6.**
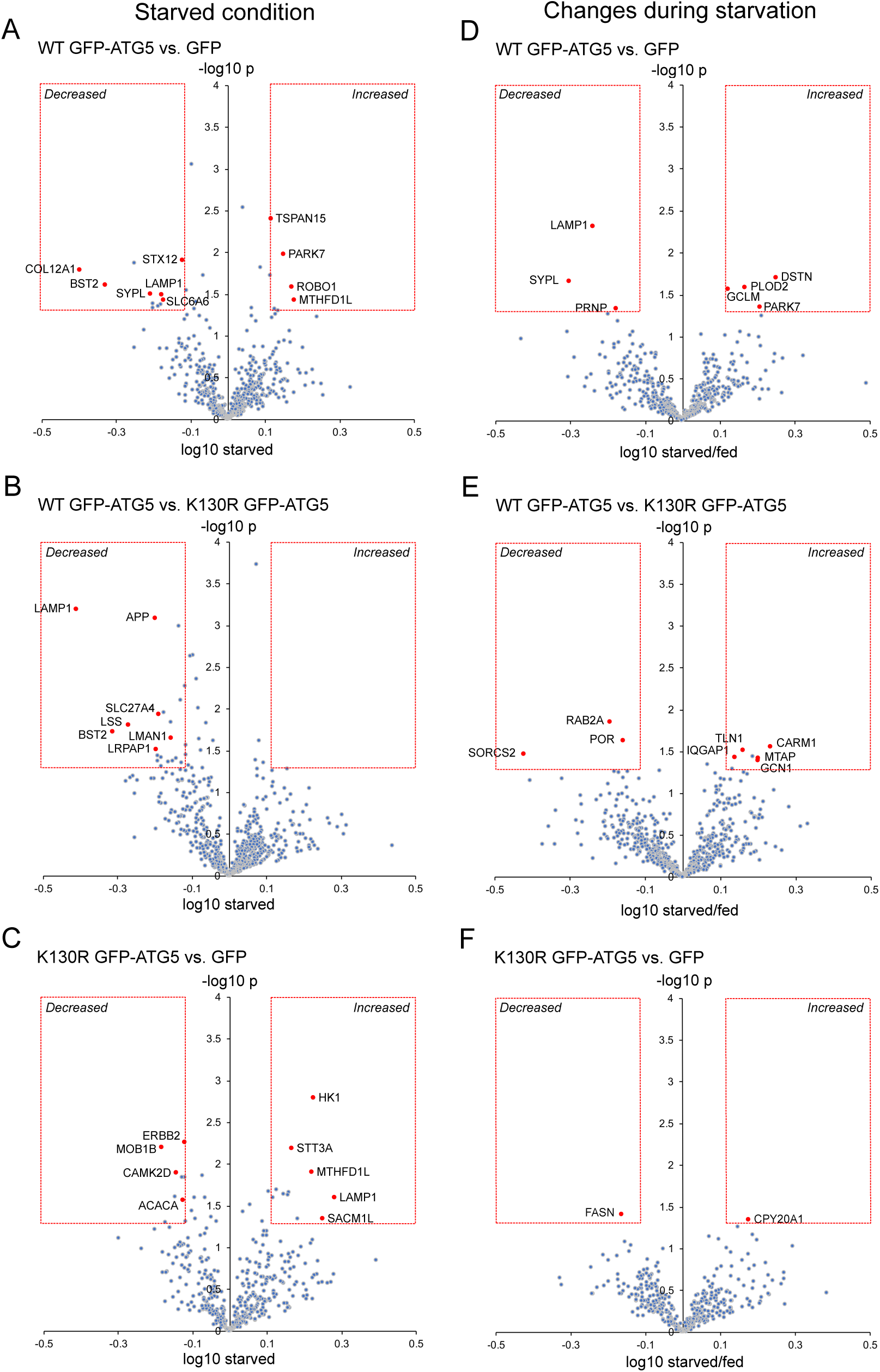
Changes in surface protein abundance during starvation in ATG5^−/−^ MEF rescued with GFP, WT GFP-ATG5 or K130R GFP-ATG5. Pairwise comparisons of surface protein abundance analysed by surface biotinylation/streptavidin pull-down and SILAC quantitative proteomics in starvation conditions. (A-C) Pairwise surface protein expression analysis following 1-hour nutrient starvation. (A) WT GFP-ATG5 vs. GFP. (B) WT GFP-ATG5 vs. K130R GFP-ATG5. (C) K130R GFP-ATG5 vs. GFP. (D-F) Analysis of changes in surface expression in fed and starved states. X-axes depict log_10_ ratio of starved/fed values to depict changes during starvation. (D) WT GFP-ATG5 vs. GFP. (E) WT GFP-ATG5 vs. K130R GFP-ATG5. (F) K130R GFP-ATG5 vs. GFP. In all examples, the red boxes indicate 1.3-fold enrichment cut-off and p<0.05. Red diamonds depict selected proteins of potential interest.

Finally, we compared the surface proteomes obtained from fed and starved WT GFP-ATG5, K130R GFP-ATG5 and GFP expressing MEFs to derive pairwise comparisons between conditions. This would reveal any proteins whose changing surface abundancies during nutrient challenge are linked to ATG12∼ATG5 conjugation and an active autophagy pathway. Comparing the WT GFP-ATG5 and GFP surface proteomes revealed relatively few proteins that changed >1.3-fold (starved/fed), these being: DSTN; PARK7; PLOD2; GCLM (all up); and PRNP; LAMP1; SYPL (all down) (**Fig. 6D; Supplemental Table 12**). Comparing WT GFP-ATG5 against K130R GFP-ATG5 surface proteomes revealed the following that significantly changed during starvation: CARM1; MTAP; GCN1; TLN1; ICGAP1 (all up); and POR; RAB2A; SORCS2 (all down) (**Fig. 6E; Supplemental Table 12**). Finally, only CPY20A1 (up) and FASN (down) were found to differ between the 2 autophagy-deficient cell-lines, K130R GFP-ATG5 and GFP (**Fig. 6F; Supplemental Table 12**). Amongst these changes, LAMP1 was again found to be decreased when comparing WT GFP-ATG5 against GFP-expressing cells (**Fig. 6D; Supplemental Table 12**). Analysis of the LAMP1 ratios under fed conditions showed very little differences between the different cell-lines (see **Supplemental Table 12**), arguing that changes in its steady state localisation only occur in response to altered cellular energy status, in the presence of an active autophagy system. LAMP1 is detected on the surface of cancer cells (e.g. breast cancer [77]) and protects NK cells from damage during degranulation [78]. It is also implicated in the release of ATP during immunogenic cell death, via packaging into autolysosomes [79]. Interestingly, mitoxantrone-induced immunogenic ATP release was found to depend on autophagy and the presence of ATG5 in this model. Whether the LAMP1- and ATG5-dependent dependent delivery of ATP sequestering autolysosomes to the plasma membrane in mitoxantrone-treated cells would correlate with altered surface LAMP1 levels [79], and whether this pathway is in any way linked to our finding that surface LAMP1 decreases on the surface of starved, autophagy-dependent cells (**Fig. 6D; Supplemental Table 12**) remains to be determined.

In summary, our investigations have revealed strong interactions between autophagosome assembly factors—specifically the ATG12∼ATG5 conjugate—and the clathrin-mediated trafficking machinery (**Fig. 2C, D**). We tested some of these interactors for their possible roles during autophagosome assembly, providing evidence that PIK3C2A and HIP1R are candidate autophagy regulators (**Figs 3, 4**). Using reconstituted ATG3 null MEFs, we revealed the association with the clathrin-mediated trafficking machinery was upstream and/or independent of ATG8 lipidation. To explore whether ATG12∼ATG5 influences clathrin-associated trafficking—we carried out surface proteomics of the MEF cell-lines in fed and starved states (**Fig. 5, 6**). Although no obvious patterns emerged in the populations of proteins having altered surface levels, a number of interesting proteins were identified whose plasma membrane distribution appeared to be influenced by the presence of ATG5 (**Fig. 5, 6**). These data can form a framework for analysis of the influence of the core autophagy machinery on intracellular trafficking in basal and stressed conditions.

## MATERIALS & METHODS

### Reagents and antibodies

Unless stated otherwise, all reagents were from Sigma: GFP-Trap beads (Chromotek, GTA-100); BafA1 (Millipore, 475984) was used at 200nM; DAPI (Life Technologies, D121490) was used at 100ng/mL; Lipofectamine 2000 (Thermo Fisher, 11668027) was used for transient transfections according to the manufacturer’s instructions; puromycin (Sigma, P8833) was used at 2*µ*g/mL. Starvation was carried out either using Hanks Balanced Salt Solution (Thermo Fisher, 14025092), or with starvation media comprising 140mM NaCl, 1mM CaCl_2_, 1mM MgCl_2_, 5mM glucose and 20mM HEPES, pH 7.4, supplemented with 1% (w/v) bovine serum albumin (BSA) [5]. Cre recombinase lentiviral particles were a gift from Prof. James Uney (Bristol).

Primary antibodies used were: anti-tubulin (Sigma, T5168); anti-ATG5 (Sigma, A0856); anti-LC3B (Sigma, L8918); anti-LC3B (Cell Signaling, 2956); anti-α-tubulin (B-5-1-2; Sigma, T6074); anti-GFP (Covance, MMS-118P); PIK3C2A (Santa Cruz, sc-40); anti-IGFR2/CI-M6PR (Abcam, ab2733); anti-TGN38 (R&D Systems, AF806); anti-SQSTM1/P62 (GeneTex, GT1478); anti-HIP1R (Abcam, ab57584); anti-ATG16L1 (MBL Life Sciences, M150-3); anti-ATG3 (MBL Life Sciences, M133-3); anti-CLTC (Sigma-Aldrich, C1860); anti-WIPI2 (AbD Serotech, MCA5780GA); anti-DNM1 (Abcam, ab13251); anti-DNM2 (Abcam, ab65556); anti-SNX9 (Santa Cruz, sc-166863). HRP-tagged secondary antibodies were from Jackson ImmunoResearch; fluorescent Alexa-tagged antibodies were from Molecular Probes.

### Cell-lines and cell culture

Unless otherwise stated, cells were grown in DMEM containing 4500mg/mL (Sigma, D5796) at 37°C with 5% CO_2_. The following cell-lines were used: ATG5 wild-type and ATG5 null MEFs, and ATG3 wild-type and ATG3-null MEFs (from Dr Sharon Tooze, CRICK Institute, London); PIK3C2A^flox/flox^ MEFs (from Prof. Yoh Takuwa, Kanazawa University, Japan, [54]); HIP1R null MEFs (from Dr Theodora Ross, UT Southwestern Medical Centre, USA, [80]; HEK293T (ATCC). ATG5- and ATG3-null MEFs stably expressing GFP, WT GFP-ATG5 or K130R GFP-ATG5 were produced by lentiviral transduction. Wild-type and K130R ATG5 cDNA (mouse sequence synthesised by Eurofins) was sub-cloned into pLVX-puro using Afe1 and BamH1 restriction sites. Lentiviruses were generated in HEK293T cells by transfection with cDNAs along with packaging vectors pMD2.G and pAX2. Lentiviral particles were collected at 48 hrs, cleared by centrifugation at 2900*g*, 10 mins, then passed through a 0.45*µ*m polyethersulfone filter to be used immediately or stored at -80°C. MEFs were transduced on 6 cm plates at ∼40% confluency by two rounds of viral addition. Cells were selected using puromycin. For SILAC, cells were grown for 8 doublings in DMEM for SILAC (Thermo, 89985), containing 4500mg/mL glucose and 4mM L-glutamine, supplemented with 100*µ*g/mL penicillin, 100*µ*g/mL streptomycin and 10% fetal bovine serum (Sigma, F0392). The following amino acids were added: “light” medium (R0/K0), Arg0 (Sigma, A6969)/Lys0 (Sigma, L8662); “medium” medium (R6/K4), Arg6 (Sigma, 643440)/Lys4 (Thermo, 88437); “heavy” medium (R10/K8), Arg10 (Silantes, 201604301)/Lys8 (Silantes, 211204302).

### GFP-Trap

For GFP-Trap immunoisolation, cells in 10cm plates were washed twice with ice cold PBS, and 1 ml of lysis buffer (10mM Tris base, 150mM NaCl, 0.5% (v/v) NP40, 1mM PMSF, 2mM MgCl_2_, protease inhibitors, pH 7.5) was added. Cells were scraped and lysates collected and incubated on ice for 20 min. Lysates were cleared by centrifugation at 20000*g* for 15 min at 4°C, and added to pre-equilibrated beads and rotated for 1 hour at 4°C. The sample was then spun at 600*g* for 2 min at 4°C to pellet beads, which were washed 3x in wash buffer (10mM Tris base; 150mM NaCl; 1mM PMSF; 2mM MgCl_2_; protease inhibitors) before they were resuspended in SDS-PAGE gel sample buffer.

### Surface biotinylation

Cells were grown to confluency in 15cm dishes. Sulfo-NHS-SS Biotin (Thermo Fisher Scientific, A39258) was prepared at 0.2mg/mL in ice-cold PBS, and cells were incubated with 5mL PBS-biotin on ice for 30 min. Cells were quenched for 10 min (50mM Tris, 100mM NaCl, pH 7.5), and then lysed in 1mL lysis buffer (PBS with 2% Triton X-100 and protease inhibitors). Cells were scraped and the lysates cleared at 20000*g* for 10 min at 4°C. The cleared lysates were then incubated with Streptavidin Sepharose (GE Healthcare, 17511301) for 30 min at 4°C under rotation. After extensive washes, samples were resuspended in SDS-PAGE gel sample buffer.

### Quantitative SILAC proteomics

Liquid chromatography coupled to tandem mass spectrometry (LC-MS/MS) analysis was carried out using an LTQ Orbitrap Velos mass spectrometer (ThermoFisher Scientific) in the Bristol University Proteomics facility. Gels were cut into 6 slices using an automated digestion unit (ProGest; Digilab UK), and a trypsin digest performed followed by peptide fractionation [nano-HPLC system (UltiMate 3000, Dionex)], ionization (ES542; Proeson) and MS/MS. MS data was acquired using Xcalibar v2.1 software (Thermo Fisher Scientific). The results were first analysed using a Sequest search against the Uniprot mouse database and then filtered to remove low confidence peptides [<5% false-discovery rate (FDR)]. Using Excel, proteins with a unique peptide number <2 were discarded. The unique peptide number displays the number of peptide sequences unique to a protein group. For the surface interactomes, data were normalised to account for any uneven incorporation of the SILAC labelled media. The median value for each condition was subtracted from each value to create a new median of 1, and Log2 of each value was obtained. A T-test was then carried out for proteins present within at least 3 of the datasets, with proteins having a fold-difference between pairwise comparisons of >1.3 and p<0.05 being considered of interest. For the starved datasets, a second analysis in which the fold change within pairwise samples was compared to the fed state was carried out (starved/fed), with fold-difference >1.3 and p<0.5 signifying interest. STRING analysis (Search Tool for the Retrieval of Interacting Genes/Proteins; http://string-db.org/) was used to identify putative protein interaction networks.

### Microscopy

Fixed and live-cell images were obtained using an Olympus IX-71 inverted microscope (60x Uplan Fluorite objective; 0.65-1.25NA, oil immersion lens) fitted with a CoolSNAP HQ CCD camera (Photometrics, AZ) driven by MetaMorph software (Molecular Devices). MetaMorph software was used to quantify puncta numbers. A TopHat morphology filter was used to score circular objects of 5 pixels (∼1um) diameter. An automated cell count was then performed to count the number of selected items. For a typical experiment, ten random fields were imaged and puncta numbers per cell in each field was counted. This was repeated three times. All other image analysis was carried out using Fiji. Confocal microscopy was carried out using a Leica SP5-AOBS confocal laser scanning microscope (63x oil immersion objective, 1.4NA; or 100x oil immersion objective, 1.4NA) attached to a Leica DM I6000 inverted epifluorescence microscope. Laser lines were: 100mW Argon (for 458, 488, 514nm excitation); 2mW Orange HeNe (594nm); and 50mW diode laser (405nm). The microscope was run using Leica LAS AF software (Leica, Germany). FRET analysis was also carried out on this microscope.

For CLEM, cells were grown on gridded 3cm imaging dishes (MatTek, P35G-1.5-14-CGR), imaged live by widefield fluorescence imaging to observe GFP-ATG5 puncta under starvation conditions, and fixed by adding glutaraldehyde to 3%. Cells were processed for transmission electron microscopy as follows: cells were washed in 0.1M sodium cacodylate, osmicated for 1h (1% osmium tetroxide in 0.1M sodium cacodylate), dehydrated by increasing ethanol concentration steps, then embedded in Epon resin. Blocks were then trimmed to the cells of interest, sectioned, then imaged using a FEI Technai 12 120kV transmission electron microscope. Images were captured using a FEI Ceta 4k x4k CCD camera.

## AUTHOR CONTRIBUTIONS

JDL designed the study and directed the research. KB carried out the experiments and interpreted data. JDL wrote the manuscript.

## ACKNOWLEDGEMENTS

The authors are grateful to Dr Kate Heesom and the University of Bristol proteomics facility. We also acknowledge the support of the Wolfson Bioimaging Facility and the BBSRC Alert 13 capital grant (BB/L014181/1); Wellcome Trust (grant no.); Natalia Jimenez-Moreno for critical reading of the manuscript.

