## Supplemental tables and figures for "The ATG5 Interactome Links Clathrin Vesicular Trafficking With The ATG8 Lipidation Machinery For Autophagosome Assembly"

#### SUPPLEMENTARY FIGURE LEGENDS

**Fig. S1 ATG5$^{-/-}$ MEFs are autophagy deficient.** Analysis of autophagy responses in wild-type and ATG5-/- MEFs by (A) immunoblotting and (B-D) immunofluorescence. Cells were starved for 1 hour in the absence or presence of BafA1. Example immunofluorescence images (B) and quantitation of WIPI2 (C) and LC3B (D) puncta are shown. Data show mean ± SEM; n=3; 10 cells per condition, per experiment. One-way ANOVA with post-hoc Tukey's test: ** $p < 0.01$; *** $p < 0.001$. Bar = 10 $\mu$m.

**Fig. S2 STRING analysis of interactions within the high confidence K130R GFP-ATG5 interactome.**

**Fig. S3 SILAC-based proteomics analysis of GFP-ATG5 interactors in the ATG3$^{-/-}$ MEF background.** (A) Immunoblots showing the autophagy response in wild-type and ATG3$^{-/-}$ MEFs subjected to starvation (1 hour) in the absence or presence of BafA1. (B) Immunoblots showing the autophagy response in ATG3$^{-/-}$ MEFs stably expressing GFP, wild-type (WT) GFP-ATG5 or K130R GFP-ATG5. Note the absence of lipidated LC3B-II. (C) STRING analysis of WT GFP-ATG5 interactors in the ATG3$^{-/-}$ MEF background. (D) STRING analysis of K130R GFP-ATG5 interactors in the ATG3$^{-/-}$ MEF background. (E) Immunoblots of GFP-TRAP pull-downs in ATG3$^{-/-}$ MEFs stably expressing GFP, WT GFP-ATG5 or K130R GFP-ATG5 showing ATG16L1, PIK3C2A and ATG5 immunoreactivity.

**Fig. S1**

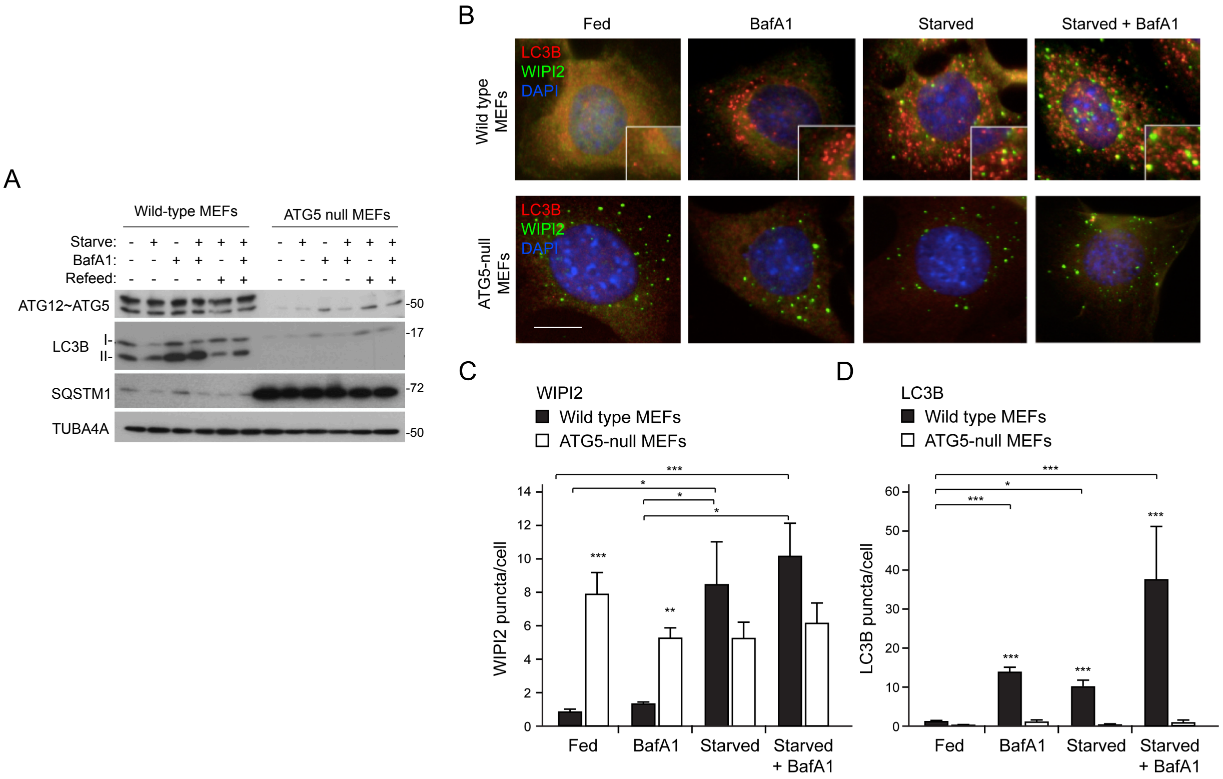
A
Wild-type MEFs
ATG5 null MEFs
Starve:
BafA1:
Refeed:
ATG12~ATG5
LC3B
I-
II-
SQSTM1
TUBA4A
-50
-17
-72
-50
B
Fed
BafA1
Starved
Starved + BafA1
Wild type MEFs
ATG5-null MEFs
LC3B
WIPI2
DAPI
C
WIPI2
Wild type MEFs
ATG5-null MEFs
WIPI2 puncta/cell
Fed
BafA1
Starved
Starved + BafA1
D
LC3B
Wild type MEFs
ATG5-null MEFs
LC3B puncta/cell
Fed
BafA1
Starved
Starved + BafA1

**Fig. S2**

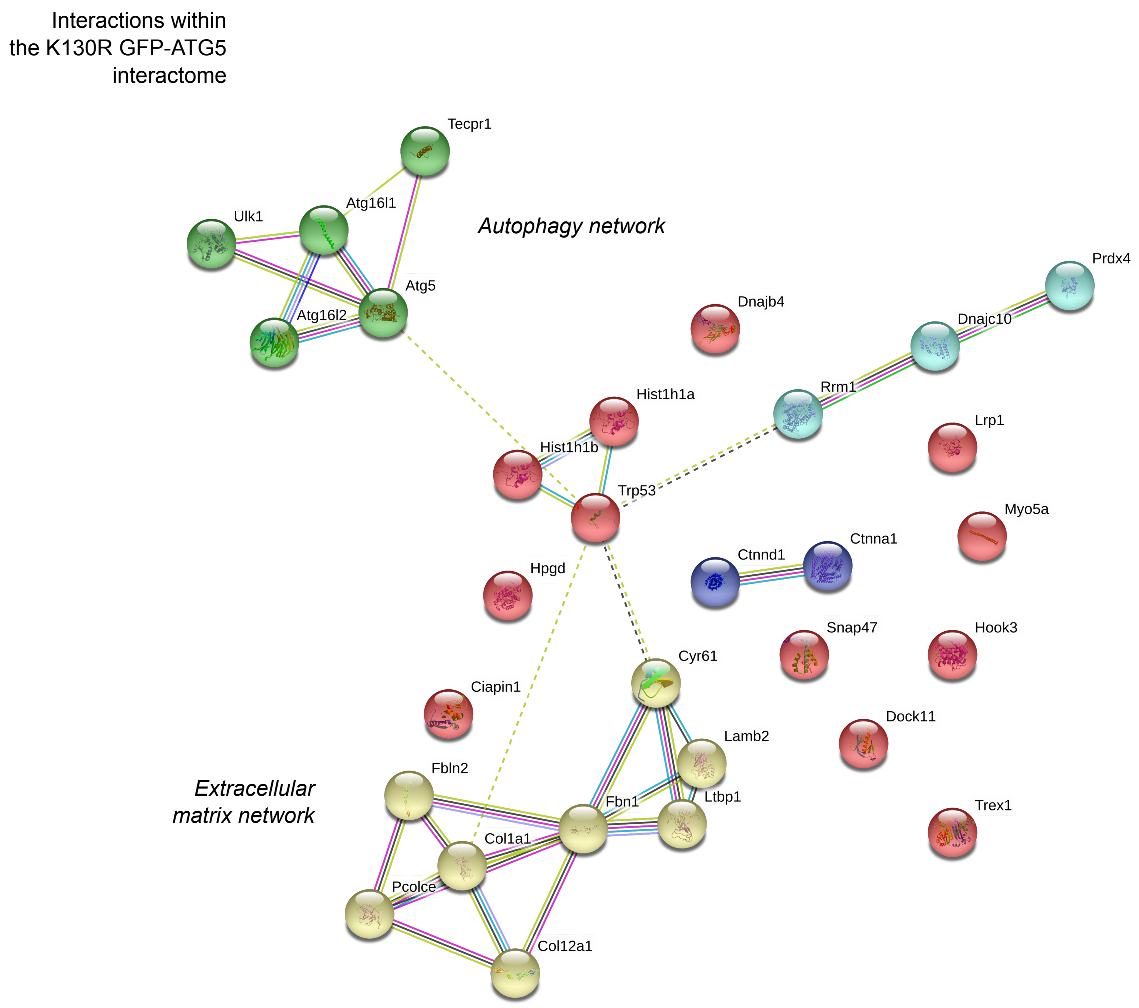
Interactions within
the K130R GFP-ATG5
interactome
Autophagy network
Extracellular
matrix network
Tecpr1
Ulk1
Atg16l1
Atg5
Atg16l2
Dnajb4
Prdx4
Dnajc10
Rrm1
Hist1h1a
Hist1h1b
Trp53
Lrp1
Myo5a
Ctnna1
Ctnnd1
Hpgd
Snap47
Hook3
Cyr61
Ciapin1
Dock11
Lamb2
Fbln2
Fbn1
Ltbp1
Trex1
Col1a1
Pcolce
Col12a1

### Fig. S3

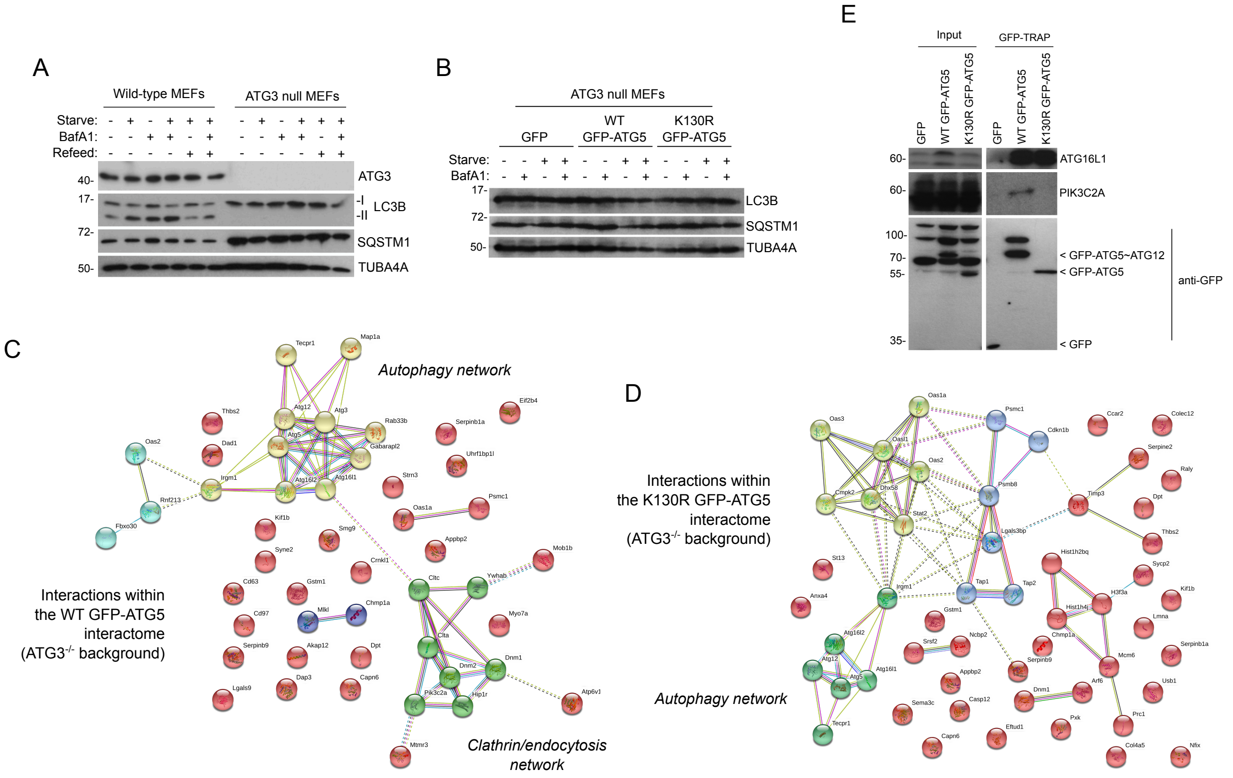

A
Wild-type MEFs
ATG3 null MEFs
Starve:
BafA1:
Refeed:
40-
17-
72-
50-
ATG3
-I
-II
LC3B
SQSTM1
TUBA4A
B
ATG3 null MEFs
GFP
WT GFP-ATG5
K130R GFP-ATG5
Starve:
BafA1:
17-
72-
50-
LC3B
SQSTM1
TUBA4A
E
Input
GFP-TRAP
GFP
WT GFP-ATG5
K130R GFP-ATG5
60-
60-
100-
70-
55-
35-
ATG16L1
PIK3C2A
< GFP-ATG5~ATG12
< GFP-ATG5
< GFP
anti-GFP
C
Autophagy network
Interactions within the WT GFP-ATG5 interactome (ATG3-/- background)
Clathrin/endocytosis network
D
Interactions within the K130R GFP-ATG5 interactome (ATG3-/- background)
Autophagy network

#### SUPPLEMENTARY TABLES

Supplementary tables 1, 4, 6 and 10 are Excel files containing complete interactions data* (**Table 1:** ATG5 interactome data; **Table 4:** ATG5 interactome data in the ATG3 null background; **Table 6:** Surface protein proteome—fed conditions—three datasheets representing WT ATG5-GFP vs. GFP, WT ATG5-GFP vs. K130R ATG5-GFP and K130R ATG5-GFP vs. GFP. P-values <0.05 are highlighted; **Table 10:** Surface protein proteome—starvation conditions—three datasheets representing WT ATG5-GFP vs. GFP, WT ATG5-GFP vs. K130R ATG5-GFP and K130R ATG5-GFP vs. GFP. P-values <0.05 are highlighted).

**These will be made available online on the journal website in the final, published version of the manuscript.*

Supplementary tables 2, 3, 5, 7, 8, 9, 11 and 12 are supplied below:

| Accession | Description | Unique peptides | WT:GFP | WT:K130R | K130R:GFP | Score |
|---|---|---|---|---|---|---|
| P60521 | GABARAPL2 | 6 | 100.000 | 100.000 | 0.010 | 71.66 |
| Q569L8 | CENPJ | 2 | 100.000 | 100.000 | 100.000 | 2.41 |
| Q3UV55 | NR1D1 | 2 | 100.000 | 100.000 | 0.040 | 0.00 |
| P39053 | DNM1 | 14 | 100.000 | 94.335 | 0.684 | 43.86 |
| O70405 | ULK1 | 9 | 100.000 | 39.869 | 100.000 | 13.22 |
| Q9JKY5 | HIP1R | 34 | 100.000 | 23.723 | 0.742 | 60.33 |
| Q3TRC8 | ARRB2 | 4 | 100.000 | 7.668 | 100.000 | 5.71 |
| Q8BW35 | Uncharacterized | 2 | 100.000 | 6.482 | 100.000 | 2.04 |
| Q8C0J2 | ATG16L1 | 13 | 100.000 | 6.478 | 27.903 | 342.83 |
| O70305 | ATXN2 | 2 | 100.000 | 5.058 | 100.000 | 1.85 |
| Q3TCY0 | RAB33B | 2 | 100.000 | 4.520 | 55.315 | 3.74 |
| D3YZ62 | MYO5A | 11 | 100.000 | 3.958 | 100.000 | 15.05 |
| Q8BTM5 | SLC2A1 | 2 | 100.000 | 3.945 | 100.000 | 0.00 |
| Q8VCC1 | HPGD | 7 | 100.000 | 3.372 | 26.545 | 32.91 |
| Q8R080 | GTSE1 | 4 | 100.000 | 3.092 | 77.471 | 5.69 |
| B1AWE0 | CLTA | 7 | 86.452 | 100.000 | 0.012 | 24.77 |
| Q9CPX6 | ATG3 | 18 | 81.825 | 100.000 | 0.010 | 324.34 |
| Q99J83 | ATG5 | 17 | 55.399 | 2.601 | 100.000 | 367.86 |
| F8VPL2 | PIK3C2A | 38 | 55.352 | 88.334 | 0.342 | 53.28 |
| F7AC41 | PUS7 | 2 | 55.194 | 2.549 | 40.840 | 0.00 |
| Q3TZU7 | SNX9 | 16 | 49.013 | 100.000 | 0.045 | 45.82 |
| Q8R3R8 | GABARAPL1 | 2 | 47.608 | 100.000 | 0.010 | 8.40 |
| Q68FD5 | CLTC | 74 | 46.149 | 100.000 | 0.114 | 600.82 |
| Q3TAB9 | ATG7 | 29 | 46.072 | 100.000 | 0.010 | 204.66 |
| Q3UIZ0 | GAK | 29 | 41.810 | 100.000 | 0.020 | 48.65 |
| Q8VD75 | HIP1 | 19 | 40.988 | 100.000 | 0.010 | 25.33 |
| G3X9G4 | DNM2 | 27 | 35.464 | 78.710 | 0.396 | 111.81 |
| Q9ESK9 | RB1CC1 | 39 | 31.434 | 100.000 | 0.011 | 48.78 |
| P62743 | AP2S1 | 3 | 28.313 | 7.944 | 2.200 | 7.62 |
| B7ZWC4 | IGF2R | 24 | 26.275 | 41.385 | 0.369 | 28.69 |
| P51863 | ATP6V0D1 | 2 | 26.249 | 2.909 | 11.648 | 4.51 |
| Q9D699 | GAS2 | 3 | 25.446 | 2.321 | 15.349 | 3.65 |
| Q9JKB1 | UCHL3 | 2 | 24.281 | 2.083 | 22.320 | 2.52 |
| G3UWG1 | GM10108 | 3 | 23.854 | 18.617 | 1.742 | 8.75 |
| Q6TXD4 | DNMBP | 8 | 23.310 | 100.000 | 0.010 | 5.19 |
| Q9DBW2 | INPP5B | 3 | 22.753 | 2.216 | 14.878 | 0.00 |
| Q99JB8 | PACSIN3 | 2 | 20.976 | 5.258 | 9.888 | 1.73 |
| Q8K382 | DENND1A | 4 | 18.387 | 100.000 | 0.010 | 5.42 |
| Q9CQY1 | ATG12 | 6 | 18.097 | 100.000 | 0.010 | 48.36 |
| G3X956 | SUPT16 | 2 | 17.497 | 3.443 | 4.406 | 0.00 |
| D3Z656 | SYNJ1 | 13 | 17.100 | 38.822 | 0.615 | 10.62 |
| E9QAT4 | SEC16A | 8 | 17.048 | 4.347 | 3.953 | 9.04 |

| | | | | | | |
|---|---|---|---|---|---|---|
| Q8CC03 | AP1G1 | 7 | 16.283 | 7.192 | 2.288 | 13.82 |
| P62192 | PSMC1 | 2 | 15.605 | 3.696 | 9.613 | 6.29 |
| P18828 | SDC1 | 3 | 15.263 | 4.064 | 0.491 | 2.08 |
| Q8BUK6 | HOOK3 | 4 | 14.959 | 3.502 | 100.000 | 7.53 |
| F6RAZ3 | ACADS | 2 | 13.991 | 14.156 | 1.000 | 5.43 |
| Q3TDG9 | STX12 | 2 | 13.990 | 100.000 | 0.088 | 0.00 |
| B2RQQ7 | CDC42BPB | 5 | 13.927 | 5.485 | 72.026 | 12.18 |
| Q3UP40 | PACSIN2 | 2 | 13.148 | 2.188 | 12.836 | 0.00 |
| E9QA74 | MYO18A | 3 | 12.559 | 10.277 | 1.222 | 1.84 |
| Q3TWV4 | AP2M1 | 14 | 12.296 | 7.057 | 1.099 | 34.76 |
| Q0PD42 | RAB13 | 2 | 11.818 | 4.300 | 2.749 | 6.11 |
| D3YZ06 | HSPB1 | 2 | 11.791 | 5.417 | 2.177 | 1.69 |
| Q9WUU7 | CTSZ | 4 | 11.683 | 2.934 | 7.891 | 6.97 |
| Q32ME1 | ATP2B4 | 2 | 11.243 | 100.000 | 0.010 | 1.82 |
| P17426 | AP2A1 | 21 | 10.909 | 15.395 | 0.609 | 83.06 |
| Q3TD51 | PICALM | 13 | 10.265 | 24.624 | 0.407 | 31.43 |
| D3Z2E3 | REPS1 | 4 | 9.675 | 47.542 | 0.273 | 3.92 |
| Q9DCE6 | DAB2 | 3 | 9.551 | 7.472 | 1.278 | 1.87 |
| P17427 | AP2A2 | 21 | 9.287 | 7.276 | 1.256 | 77.01 |
| P35585 | AP1M1 | 11 | 9.042 | 14.268 | 0.973 | 13.52 |
| Q5SVG5 | AP1B1 | 6 | 8.141 | 6.996 | 1.164 | 64.85 |
| Q9DBG3 | AP2B1 | 11 | 8.085 | 7.838 | 0.987 | 81.04 |
| Q3UHX2 | PDAP1 | 3 | 7.690 | 6.013 | 1.279 | 6.80 |
| Q6A009 | LTN1 | 3 | 7.541 | 2.226 | 3.904 | 3.31 |
| E9PYE2 | TYW3 | 2 | 7.280 | 2.288 | 3.182 | 2.38 |
| G3X920 | ARMC8 | 3 | 6.708 | 10.846 | 0.618 | 2.68 |
| D3Z7E5 | GSK3A | 4 | 6.089 | 2.657 | 2.037 | 2.33 |
| Q6PIU9 | FLJ45252 HOMOLOG | 2 | 6.048 | 68.229 | 0.089 | 3.72 |
| Q920I9 | WDR7 | 2 | 5.877 | 6.022 | 0.463 | 1.93 |
| O54781 | SRPK2 | 4 | 5.847 | 4.766 | 0.583 | 8.29 |
| Q3V471 | LGALS3 | 2 | 5.752 | 3.893 | 1.477 | 1.78 |
| A2AUM2 | EIF2AK4 | 2 | 5.664 | 100.000 | 0.010 | 0.00 |
| Q80SW1 | AHCYL1 | 10 | 5.520 | 5.205 | 0.903 | 25.09 |
| H9KUZ8 | CEP41 | 2 | 5.499 | 2.720 | 2.021 | 1.89 |
| Q01320 | TOP2A | 3 | 5.434 | 2.332 | 2.330 | 1.97 |
| Q9DB05 | NAPA | 8 | 5.344 | 2.545 | 1.331 | 16.51 |
| P97390 | VPS45 | 4 | 5.330 | 2.621 | 23.458 | 3.58 |
| Q9CW46 | RAVER1 | 5 | 5.143 | 3.066 | 0.104 | 6.47 |
| A0A0A0MQN9 | TTC28 | 4 | 4.975 | 2.872 | 1.732 | 1.69 |
| Q3UAZ7 | HMGB2 | 2 | 4.951 | 3.456 | 1.433 | 3.78 |
| Q8BXV2 | BRI3BP | 2 | 4.937 | 3.257 | 1.516 | 3.38 |
| D3Z6W1 | MAP6 | 2 | 4.780 | 5.552 | 0.861 | 6.11 |
| D3Z7S0 | GET4 | 2 | 4.732 | 2.519 | 1.879 | 4.05 |
| Q9R061 | NUBP2 | 2 | 4.694 | 2.776 | 1.691 | 13.83 |
| Q8VHC3 | SELM | 2 | 4.661 | 2.910 | | 2.11 |

| | | | | | | |
|---|---|---|---|---|---|---|
| Q3TYL9 | CNP | 4 | 4.525 | 2.537 | 1.768 | 12.95 |
| O35682 | MYADM | 2 | 4.394 | 3.044 | 1.443 | 2.03 |
| Q80ZU1 | DCAF7 | 2 | 4.384 | 15.638 | 0.121 | 1.84 |
| P70271 | PDLIM4 | 5 | 4.363 | 3.587 | 1.085 | 5.58 |
| Q640N1 | AEBP1 | 2 | 4.325 | 3.305 | 1.309 | 1.61 |
| B7ZMQ9 | INPP4A | 2 | 4.317 | 3.778 | 0.857 | 1.72 |
| Q08091 | CNN1 | 8 | 4.278 | 7.478 | 0.551 | 29.74 |
| Q9EPK6 | SIL1 | 3 | 4.245 | 100.000 | 0.010 | 5.30 |
| Q9QYJ3 | DNAJB1 | 4 | 4.194 | 2.047 | 2.049 | 5.17 |
| Q3UHJ0 | AAK1 | 7 | 4.169 | 100.000 | 0.010 | 6.44 |
| A2AFQ9 | GEMIN5 | 8 | 4.099 | 2.086 | 2.406 | 8.94 |
| Q9QZA0 | CA5B | 2 | 4.068 | 11.774 | 0.284 | 0.00 |
| F8VQE9 | AGAP3 | 3 | 4.029 | 2.554 | 1.577 | 3.77 |
| P56389 | CDA | 2 | 3.948 | 3.280 | 1.204 | 2.89 |
| Q8C871 | SERPINB8 | 3 | 3.904 | 5.142 | 0.746 | 6.37 |
| F8WHM5 | GLG1 | 8 | 3.822 | 5.653 | 0.660 | 14.36 |
| Q6SLK2 | WNK1 | 4 | 3.796 | 3.984 | 0.953 | 0.00 |
| Q80XP8 | FAM76B | 2 | 3.716 | 2.580 | 1.440 | 4.02 |
| Q3UGK7 | WASL | 3 | 3.639 | 2.156 | 1.688 | 1.98 |
| F8VPN4 | AGL | 6 | 3.600 | 100.000 | 0.010 | 3.55 |
| Q3UXP1 | PHF6 | 2 | 3.471 | 2.400 | 1.446 | 0.00 |
| P62331 | ARF6 | 2 | 3.436 | 100.000 | 0.010 | 0.00 |
| Q8C078 | CAMKK2 | 4 | 3.368 | 14.148 | 0.086 | 1.71 |
| Q9CQV8 | YWHAB | 3 | 3.315 | 2.258 | 1.441 | 42.77 |
| Q08288 | LYAR | 3 | 3.292 | 2.756 | 1.194 | 1.70 |
| A0A0B4J1E5 | UCK2 | 2 | 3.289 | 11.609 | 0.088 | 5.75 |
| Q3TDW6 | APBB2 | 2 | 3.254 | 2.004 | 1.624 | 0.00 |
| F6XC25 | CC2D1B | 3 | 3.211 | 3.077 | 1.044 | 7.26 |
| Q6PAR5 | GAPVD1 | 2 | 3.208 | 4.789 | 0.670 | 0.00 |
| P80314 | CCT2 | 15 | 3.157 | 2.418 | 1.113 | 77.57 |
| Q80VY1 | HZGJ-LIKE | 2 | 3.108 | 6.041 | 0.893 | 6.05 |
| O08547 | SEC22B | 4 | 3.101 | 2.725 | 1.362 | 10.18 |
| Q8BTU6 | EIF4A2 | 4 | 3.062 | 22.427 | 0.082 | 45.68 |
| B1AZ15 | COBLL1 | 3 | 3.046 | 9.333 | 0.326 | 1.64 |
| Q61879 | MYH10 | 7 | 3.032 | 3.999 | 0.738 | 43.23 |
| Q69ZG0 | MKIAA1574 | 3 | 2.962 | 2.526 | 1.419 | 0.00 |
| Q0VF62 | BCAS3 | 4 | 2.941 | 3.530 | 1.144 | 1.60 |
| Q6P1G0 | HEATR6 | 3 | 2.923 | 2.417 | 1.051 | 1.86 |
| Q9WVS5 | CCTQ | 25 | 2.889 | 2.188 | 1.221 | 113.20 |
| Q60960 | KPNA1 | 3 | 2.870 | 2.013 | 1.807 | 39.65 |
| Q5JC28 | EPS15 | 4 | 2.861 | 100.000 | 0.010 | 1.63 |
| Q9D1M0 | SEC13 | 5 | 2.849 | 2.067 | 1.151 | 13.02 |
| Q8VDP2 | CXorf56 | 4 | 2.845 | 2.781 | 0.859 | 4.16 |
| D3YZU6 | HAGHL | 3 | 2.838 | 13.021 | 0.153 | 5.98 |
| Q3UFY8 | TRMT10C | 2 | 2.830 | 28.824 | 0.055 | 2.03 |

| | | | | | | |
|---|---|---|---|---|---|---|
| P98192 | GNPAT | 3 | 2.801 | 12.088 | 0.197 | 3.49 |
| Q9JJ62 | TOP3B | 3 | 2.759 | 5.777 | 0.478 | 0.00 |
| G3X8U8 | PARG | 5 | 2.716 | 2.194 | 1.011 | 7.56 |
| Q9WUQ2 | PREB | 5 | 2.712 | 2.870 | 1.057 | 13.31 |
| Q6P9Q4 | FHOD1 | 5 | 2.705 | 2.896 | 0.696 | 7.81 |
| A2ALF0 | DNAJC8 | 3 | 2.701 | 3.215 | 0.840 | 6.57 |
| Q3TXV7 | HEXA | 2 | 2.693 | 2.561 | 1.052 | 0.00 |
| A1E2B8 | Inducible HSP70 | 10 | 2.683 | 2.243 | 1.226 | 89.40 |
| Q8BI72 | CDKN2AIP | 5 | 2.682 | 2.175 | 1.345 | 20.98 |
| Q3TUY5 | 5730455P16RIK | 3 | 2.680 | 2.366 | 1.133 | 3.77 |
| G5E896 | EDC4 | 11 | 2.680 | 2.640 | 0.845 | 15.38 |
| P27612 | PLAA | 9 | 2.672 | 2.632 | 0.534 | 11.59 |
| Q3U3U3 | GPC1 | 2 | 2.661 | 100.000 | 0.010 | 1.76 |
| E9QMH7 | IKBIP | 2 | 2.647 | 2.629 | 0.613 | 7.68 |
| Q9D6K8 | FUNDC2 | 2 | 2.644 | 2.536 | 1.042 | 0.00 |
| Q4VA53 | PDS5B | 2 | 2.633 | 100.000 | 0.010 | 4.23 |
| Q4V9X0 | PPP3CA | 2 | 2.624 | 100.000 | 0.010 | 2.30 |
| Q8BIJ7 | RUFY1 | 4 | 2.619 | 6.977 | 0.010 | 3.37 |
| E0CY23 | HSPA4L | 10 | 2.584 | 2.815 | 1.021 | 20.61 |
| A2BE92 | SET | 2 | 2.578 | 2.391 | 1.078 | 3.34 |
| A2A6U3 | SEPT9 | 11 | 2.570 | 2.013 | 1.393 | 22.47 |
| Q0KL02 | TRIO | 9 | 2.567 | 4.894 | 0.496 | 3.93 |
| D3Z3J6 | PAIP1 | 3 | 2.548 | 100.000 | 0.010 | 5.73 |
| Q9D289 | TRAPPC6B | 2 | 2.532 | 2.238 | 0.959 | 0.00 |
| Q8K297 | COLGALT1 | 7 | 2.514 | 2.002 | 1.310 | 9.60 |
| P80317 | CCT6A | 19 | 2.494 | 2.197 | 1.125 | 54.20 |
| E9QNY8 | SACS | 5 | 2.493 | 2.686 | 0.928 | 3.81 |
| Q99J95 | CDK9 | 8 | 2.490 | 2.384 | 1.337 | 35.68 |
| B2RWX4 | RINT1 | 2 | 2.475 | 3.410 | 0.726 | 0.00 |
| A2AWA9 | RABGAP1 | 2 | 2.473 | 3.809 | 0.649 | 6.03 |
| Q924H5 | RAD51C | 3 | 2.462 | 2.632 | 0.935 | 7.52 |
| D3Z198 | MRPS17 | 2 | 2.458 | 4.280 | 0.584 | 0.00 |
| H3BKG0 | CAV1 | 2 | 2.455 | 2.116 | 1.308 | 2.08 |
| Q3TX72 | FKBP5 | 2 | 2.445 | 3.308 | 0.739 | 4.11 |
| O89032 | SH3PXD2A | 2 | 2.443 | 14.036 | 0.010 | 4.57 |
| D3YU01 | PLEKHA1 | 3 | 2.431 | 2.002 | 11.216 | 3.39 |
| F6V084 | TMX1 | 2 | 2.425 | 2.475 | 0.623 | 13.55 |
| Q8C9Y7 | ECT2 | 2 | 2.422 | 2.495 | 0.951 | 0.00 |
| F6THK7 | IRAK1 | 2 | 2.419 | 2.107 | 1.148 | 7.49 |
| Q3TNH0 | TMPO | 4 | 2.418 | 2.306 | 0.981 | 2.04 |
| Q3TJ21 | PYCR2 | 6 | 2.394 | 2.872 | 1.301 | 18.20 |
| Q3TV90 | LUC7L | 2 | 2.368 | 4.244 | 0.329 | 8.87 |
| D3Z3F8 | SPG20 | 6 | 2.351 | 2.814 | 0.731 | 11.26 |
| Q8BI84 | MIA3 | 5 | 2.333 | 6.098 | 0.076 | 5.18 |
| K7Q751 | PTK2 | 6 | 2.330 | 2.087 | 0.897 | 10.55 |

| | | | | | | |
|---|---|---|---|---|---|---|
| Q3TM37 | SNW1 | 2 | 2.319 | 2.791 | 0.831 | 3.81 |
| P02469 | LAMB1 | 5 | 2.311 | 2.192 | 0.632 | 13.56 |
| Q8VDM6 | HNRNPUL1 | 5 | 2.287 | 6.112 | 0.498 | 3.49 |
| Q3U896 | MTMR9 | 2 | 2.285 | 100.000 | 0.010 | 4.34 |
| Q8BVQ0 | WDR61 | 2 | 2.278 | 3.697 | 0.616 | 2.20 |
| Q8R216 | SIRT4 | 3 | 2.278 | 2.151 | 0.908 | 12.24 |
| P80315 | CCT4 | 17 | 2.271 | 2.062 | 1.016 | 106.00 |
| Q80ZM5 | H1FX | 3 | 2.263 | 2.464 | 1.021 | 12.98 |
| B2RY04 | DOCK5 | 2 | 2.255 | 2.625 | 0.859 | 4.01 |
| V9GXD1 | CCDC132 | 2 | 2.253 | 2.427 | 0.928 | 4.19 |
| Q8BK72 | MRPS27 | 3 | 2.239 | 2.655 | 0.843 | 7.70 |
| P63101 | YWHAZ | 8 | 2.232 | 2.298 | 0.883 | 53.22 |
| Q78HU3 | MVB12A | 3 | 2.223 | 2.978 | 0.812 | 5.42 |
| P35831 | PTPN12 | 4 | 2.206 | 2.942 | 0.749 | 5.78 |
| P11983 | TCP1 | 21 | 2.205 | 2.133 | 1.007 | 79.23 |
| A2AAY5 | SH3PXD2B | 2 | 2.188 | 2.364 | 0.926 | 2.12 |
| O08614 | UTRN | 16 | 2.174 | 3.039 | 0.618 | 29.15 |
| B2RY79 | DOCK9 | 10 | 2.157 | 26.680 | 0.050 | 11.63 |
| Q99LE7 | PXN | 2 | 2.146 | 2.963 | 0.652 | 1.63 |
| Q3TYJ0 | STUB1 | 2 | 2.143 | 4.205 | 0.631 | 6.22 |
| Q99PB4 | MAGED2 | 2 | 2.141 | 2.985 | 0.787 | 3.48 |
| Q6KAP1 | MFLJ00246 | 2 | 2.140 | 2.530 | 0.846 | 20.25 |
| P80316 | CCT5 | 19 | 2.138 | 2.273 | 0.928 | 51.65 |
| Q3TSX8 | TOMM70A | 5 | 2.131 | 5.047 | 0.413 | 11.94 |
| Q9CZP0 | UFSP1 | 2 | 2.106 | 2.282 | 0.923 | 0.00 |
| Q9D786 | HAUS5 | 2 | 2.101 | 2.311 | 0.904 | 4.11 |
| P80318 | CCT3 | 29 | 2.089 | 2.023 | 1.061 | 120.84 |
| P50544 | ACADVL | 18 | 2.075 | 2.100 | 0.841 | 66.17 |
| P69566 | RANBP9 | 2 | 2.060 | 2.370 | 0.869 | 2.06 |
| O70481 | UBR1 | 3 | 2.057 | 2.016 | 0.824 | 4.02 |
| D3Z030 | LRRC16A | 10 | 2.055 | 3.274 | 0.648 | 11.65 |
| Q8C650 | SEPT10 | 2 | 2.049 | 2.633 | 0.778 | 20.08 |
| Q3ULN6 | SCCPDH | 6 | 2.046 | 8.172 | 0.250 | 9.08 |
| Q9D0I4 | STX17 | 3 | 2.041 | 3.665 | 0.638 | 2.73 |
| Q3TNW7 | TBC1D1 | 2 | 2.033 | 3.118 | 0.652 | 3.46 |
| Q9JJV2 | PFN2 | 5 | 2.028 | 2.966 | 0.705 | 46.80 |
| A2AIW9 | PMPCA | 7 | 2.024 | 5.994 | 0.313 | 16.31 |
| E9Q3L2 | PI4KA | 4 | 2.022 | 2.966 | 1.448 | 1.98 |
| P70451 | FER | 3 | 2.013 | 100.000 | 0.010 | 5.79 |
| Q6ZPE2 | SBF1 | 6 | 2.013 | 2.728 | 1.496 | 5.65 |
| P51660 | HSD17B4 | 13 | 2.001 | 2.271 | 0.943 | 63.20 |

**Supplemental Table 2:** The wild-type GFP-ATG5 interactome. Putative interactors represented by 2 or more unique peptides ranked in order of: (i) wild type GFP-ATG5:GFP interactors (>2-fold enrichment); (ii) wild-type:mutant GFP-ATG5 interactors (>2-fold enrichment); (iii) score. Autophagy proteins are highlighted green; membrane trafficking proteins are highlighted yellow.

| Accession | DESCRIPTION | Unique peptides | K130R:GFP | K130R:WT | WT:GFP | Score |
|---|---|---|---|---|---|---|
| Q9DC23 | DNAJC10 | 16 | 100.000 | 100.000 | 1.000 | 33.55 |
| Q61292 | LAMB2 | 9 | 100.000 | 100.000 | 1.000 | 14.88 |
| Q3UGP0 | DCLK1 | 5 | 100.000 | 100.000 | 1.000 | 11.07 |
| Q8CG19 | LTBP1 | 8 | 100.000 | 100.000 | 1.000 | 10.18 |
| Q3U5K8 | IFIT1 | 4 | 100.000 | 100.000 | 1.000 | 9.13 |
| Q9EQH2 | ERAP1 | 7 | 100.000 | 100.000 | 1.000 | 7.36 |
| Q3UUU0 | EMILIN1 | 6 | 100.000 | 100.000 | 1.000 | 6.73 |
| Q91VM5 | RBMXL1 | 3 | 100.000 | 100.000 | 1.000 | 6.00 |
| Q62356 | FSTL1 | 5 | 100.000 | 100.000 | 1.000 | 5.98 |
| Q8K173 | COL3A1 | 3 | 100.000 | 100.000 | 1.000 | 4.26 |
| D3Z1C5 | LDB1 | 2 | 100.000 | 100.000 | 1.000 | 3.71 |
| Q542W1 | IL1RN | 2 | 100.000 | 100.000 | 1.000 | 3.12 |
| Q9D787 | PPIL2 | 2 | 100.000 | 100.000 | 1.000 | 2.72 |
| F7C279 | BNIP1 | 2 | 100.000 | 100.000 | 1.000 | 2.58 |
| Q8VDC1 | FYCO1 | 2 | 100.000 | 100.000 | 1.000 | 2.57 |
| H7BWY5 | PARP9 | 2 | 100.000 | 100.000 | 1.000 | 2.17 |
| Q3UPI9 | POLE | 3 | 100.000 | 100.000 | 1.000 | 2.01 |
| E9Q7G0 | NUMA1 | 3 | 100.000 | 100.000 | 1.000 | 1.98 |
| F6Z1C2 | EFEMP2 | 2 | 100.000 | 100.000 | 1.000 | 1.83 |
| Q5NCU4 | SPARC | 2 | 100.000 | 100.000 | 1.000 | 1.74 |
| Q14AX3 | KCTD12 | 2 | 100.000 | 100.000 | 1.000 | 1.70 |
| P83870 | PHF5A | 3 | 100.000 | 100.000 | 1.000 | 1.62 |
| D3YUE0 | TREX1 | 3 | 100.000 | 22.219 | 1.000 | 7.52 |
| Q6AXC6 | DDX11 | 2 | 100.000 | 15.577 | 10.000 | 1.61 |
| Q9D6T0 | NOSIP | 3 | 100.000 | 14.439 | 1.000 | 2.14 |
| P11087 | COL1A1 | 11 | 100.000 | 14.097 | 1.949 | 26.73 |
| B2RXC8 | PPP2R3A | 2 | 100.000 | 13.596 | 1.000 | 1.69 |
| A0A087WS27 | FAM46A | 2 | 100.000 | 10.468 | 1.000 | 5.55 |
| P51949 | MNAT1 | 3 | 100.000 | 10.101 | 10.000 | 1.63 |
| B1B0C7 | HSPG2 | 8 | 100.000 | 2.538 | 1.000 | 1.65 |
| Q9CQW9 | IFITM3 | 2 | 77.242 | 77.242 | 1.000 | 1.68 |
| Q3TGL4 | FBLN2 | 19 | 76.850 | 17.559 | 1.341 | 53.74 |
| Q3UGQ1 | TRP53 | 10 | 75.272 | 37.923 | 1.000 | 15.36 |
| E9PX70 | COL12A1 | 24 | 72.752 | 11.089 | 1.645 | 30.54 |
| Q9Z315 | SART1 | 2 | 72.471 | 72.471 | 2.059 | 1.82 |
| Q05BJ7 | OAS1G | 2 | 42.294 | 32.177 | 1.314 | 4.71 |
| Q8BG46 | ANKRD10 | 2 | 35.353 | 16.644 | 2.124 | 2.86 |
| Q9JHT5 | AMMECR1 | 3 | 34.905 | 3.667 | 15.534 | 6.58 |
| Q922M3 | KCTD10 | 2 | 33.480 | 33.480 | 0.830 | 3.66 |
| O08807 | PRDX4 | 4 | 28.253 | 4.080 | 1.326 | 59.69 |
| Z4YJU8 | GOLGA2 | 2 | 25.105 | 2.116 | 20.770 | 1.88 |
| P09055 | ITGB1 | 6 | 24.298 | 5.649 | 3.946 | 6.13 |
| Q3TAP5 | TRA2A | 2 | 22.381 | 14.767 | 1.516 | 2.06 |

| | | | | | | |
|---|---|---|---|---|---|---|
| A0A087WPL1 | PDLIM2 | 2 | 22.370 | 14.826 | 1.509 | 4.53 |
| Q91ZX7 | LRP1 | 38 | 22.261 | 5.078 | 2.229 | 56.74 |
| Q9WVH9 | FBLN5 | 2 | 18.774 | 16.175 | 0.010 | 2.24 |
| Q8WTY4 | CIAPIN1 | 6 | 17.542 | 5.418 | 2.429 | 16.29 |
| Q3UY34 | C12orf43 | 2 | 16.114 | 7.697 | 2.094 | 5.58 |
| Q5FWI3 | TMEM2 | 4 | 14.656 | 12.720 | 1.152 | 4.01 |
| B7ZWI2 | 1110005A23RIK | 4 | 14.367 | 6.662 | 1.800 | 6.85 |
| P07742 | RRM1 | 13 | 14.111 | 9.287 | 1.162 | 27.98 |
| Q9CT37 | HNRNPR | 2 | 13.079 | 10.063 | 1.300 | 15.09 |
| Q80U95 | UBE3C | 3 | 12.107 | 7.684 | 1.576 | 4.27 |
| Q8BJY1 | PSMD5 | 2 | 12.032 | 9.537 | 1.262 | 1.79 |
| A2RSY6 | TRMT1L | 2 | 11.434 | 11.624 | 0.984 | 1.65 |
| Q6ZQJ9 | NCAPH | 5 | 11.415 | 8.735 | 1.708 | 6.92 |
| A2A9P2 | NADK | 3 | 11.253 | 2.019 | 6.119 | 6.65 |
| Q3V028 | CYLD | 2 | 11.138 | 7.927 | 1.405 | 1.78 |
| Q78E06 | EIF2AK2 | 2 | 10.817 | 12.948 | 0.835 | 4.37 |
| Q921Q3 | ALG1 | 2 | 10.253 | 8.969 | 1.143 | 1.92 |
| P04184 | TK1 | 4 | 10.173 | 2.777 | 5.029 | 4.64 |
| Q61398 | PCOLCE | 8 | 9.728 | 3.787 | 1.688 | 18.09 |
| B9EKC1 | PASK | 4 | 9.400 | 7.213 | 1.811 | 1.60 |
| P18406 | CYR61 | 6 | 8.731 | 8.030 | 2.033 | 8.10 |
| Q8VHC5 | CABP4 | 2 | 8.547 | 5.743 | 1.488 | 2.13 |
| A2AKI5 | ITGAV | 2 | 7.607 | 3.034 | 2.507 | 1.96 |
| Q3TDD1 | LDLR | 6 | 6.896 | 4.024 | 1.762 | 9.26 |
| Q6GU23 | STAT3 | 17 | 6.773 | 2.691 | 1.591 | 26.08 |
| Q9R1P1 | PSMB3 | 3 | 6.612 | 5.801 | 1.317 | 3.52 |
| Q3TC83 | NLE1 | 3 | 6.456 | 2.828 | 2.283 | 1.86 |
| P24288 | BCAT1 | 7 | 6.424 | 2.326 | 1.777 | 15.04 |
| F8WIE5 | HECTD1 | 2 | 6.239 | 15.351 | 0.010 | 1.64 |
| P48759 | PTX3 | 7 | 5.720 | 6.171 | 1.000 | 11.94 |
| Q99J10 | CTU1 | 4 | 5.674 | 13.305 | 0.141 | 6.21 |
| G3UXB4 | CTU2 | 5 | 5.666 | 3.440 | 1.949 | 18.45 |
| E9Q197 | GLOD4 | 9 | 5.513 | 1.972 | 1.965 | 32.82 |
| Q3U8R9 | TXNL1 | 8 | 5.453 | 2.482 | 2.580 | 25.44 |
| Q9WVQ5 | APIP | 7 | 5.316 | 2.323 | 1.229 | 18.86 |
| F6WM75 | CCDC167 | 2 | 5.051 | 2.023 | 2.497 | 4.00 |
| Q8VBZ3 | CLPTM1 | 2 | 4.678 | 8.796 | 0.532 | 4.16 |
| Q3THQ5 | STIP1 | 11 | 4.565 | 2.055 | 2.190 | 19.95 |
| G3UYB1 | CHEK1 | 5 | 4.492 | 3.640 | 1.155 | 12.03 |
| Q7TMW1 | RANGAP1 | 13 | 4.439 | 2.355 | 1.507 | 27.07 |
| B2RUG7 | ZFR | 5 | 4.218 | 11.632 | 1.766 | 5.14 |
| P70302 | STIM1 | 2 | 4.212 | 3.387 | 1.244 | 1.61 |
| Q8C050 | RPS6KA5 | 2 | 4.199 | 2.645 | 1.588 | 1.62 |
| E9Q7B0 | P4HA1 | 5 | 3.830 | 4.598 | 0.936 | 6.62 |
| Q8BH83 | ANKRD9 | 6 | 3.643 | 2.552 | 1.193 | 6.40 |

| | | | | | | |
|---|---|---|---|---|---|---|
| Q3U4J9 | CDK6 | 8 | 3.604 | 2.140 | 1.525 | 48.71 |
| Q3UKJ7 | SMU1 | 3 | 3.460 | 2.223 | 1.557 | 5.68 |
| P30285 | CDK4 | 8 | 3.331 | 2.325 | 1.205 | 49.91 |
| A2AU61 | RALY | 4 | 3.282 | 2.019 | 1.916 | 6.97 |
| Q8C872 | TFRC | 5 | 3.139 | 3.098 | 1.077 | 10.53 |
| Q8BGR9 | UBLCP1 | 2 | 3.122 | 2.246 | 1.627 | 1.72 |
| B1AQR8 | LGALS9 | 5 | 2.479 | 8.303 | 0.497 | 22.60 |
| G3X8P6 | TXNRD3 | 5 | 2.414 | 2.137 | 0.140 | 10.77 |
| Q9JMH6 | TXNRD1 | 7 | 2.407 | 2.927 | 0.998 | 17.86 |
| B2RQ68 | LUZP1 | 3 | 2.389 | 11.894 | 1.300 | 7.79 |
| Q9R1S8 | CAPN7 | 3 | 2.302 | 1.992 | 1.254 | 1.90 |
| Q8BHK9 | ERCC6L | 3 | 2.277 | 2.413 | 1.947 | 3.63 |
| E9Q555 | RNF213 | 59 | 2.235 | 2.579 | 0.941 | 80.39 |
| Q6PE06 | KLHDC4 | 4 | 2.176 | 2.884 | 0.581 | 4.43 |
| Q3TC83 | NLE1 | 3 | 6.456 | 2.828 | 2.283 | 1.86 |
| P04184 | TK1 | 4 | 10.173 | 2.777 | 5.029 | 4.64 |
| Q6GU23 | STAT3 | 17 | 6.773 | 2.691 | 1.591 | 26.08 |
| Q8C050 | RPS6KA5 | 2 | 4.199 | 2.645 | 1.588 | 1.62 |
| E9Q555 | RNF213 | 59 | 2.235 | 2.579 | 0.941 | 80.39 |
| Q8BH83 | ANKRD9 | 6 | 3.643 | 2.552 | 1.193 | 6.40 |
| B1B0C7 | HSPG2 | 8 | 100.000 | 2.538 | 1.000 | 1.65 |
| Q3U8R9 | TXNL1 | 8 | 5.453 | 2.482 | 2.580 | 25.44 |
| F6QN75 | LMAN2L | 2 | 3.219 | 2.472 | 1.302 | 0.00 |
| Q8BHK9 | ERCC6L | 3 | 2.277 | 2.413 | 1.947 | 3.63 |
| Q7TMW1 | RANGAP1 | 13 | 4.439 | 2.355 | 1.507 | 27.07 |
| P24288 | BCAT1 | 7 | 6.424 | 2.326 | 1.777 | 15.04 |
| P30285 | CDK4 | 8 | 3.331 | 2.325 | 1.205 | 49.91 |
| Q9WVQ5 | APIP | 7 | 5.316 | 2.323 | 1.229 | 18.86 |
| Q8BGR9 | UBLCP1 | 2 | 3.122 | 2.246 | 1.627 | 1.72 |
| A0A0A6YWN9 | USP19 | 4 | 3.399 | 2.239 | 1.000 | 0.00 |
| Q3UKJ7 | SMU1 | 3 | 3.460 | 2.223 | 1.557 | 5.68 |
| Q3U4J9 | CDK6 | 8 | 3.604 | 2.140 | 1.525 | 48.71 |
| G3X8P6 | TXNRD3 | 5 | 2.414 | 2.137 | 0.140 | 10.77 |
| Z4YJU8 | GOLGA2 | 2 | 25.105 | 2.116 | 20.770 | 1.88 |
| Q3THQ5 | STIP1 | 11 | 4.565 | 2.055 | 2.190 | 19.95 |
| F2WWK5 | FADS2 | 2 | 10.744 | 2.055 | 4.349 | 0.00 |
| F6WM75 | CCDC167 | 2 | 5.051 | 2.023 | 2.497 | 4.00 |
| A2A9P2 | NADK | 3 | 11.253 | 2.019 | 6.119 | 6.65 |
| A2AU61 | RALY | 4 | 3.282 | 2.019 | 1.916 | 6.97 |

**Supplemental Table 3:** The K130R GFP-ATG5 interactome. Putative interactors represented by 2 or more unique peptides ranked in order of: (i) K130R GFP-ATG5:GFP interactors (>2-fold enrichment); (ii) K130R GFP-ATG5:WT GFP-ATG5 interactors (>2-fold enrichment); (iii) score. Autophagy proteins are highlighted green; membrane trafficking proteins are highlighted yellow.

| Accession | Description | Unique peptides | WT:GFP | WT:K130R | K130R:GFP | Score |
|---|---|---|---|---|---|---|
| Q3TDQ5 | ATG16L1 | 31 | 100.000 | 6.120 | 32.964 | 432.04 |
| Q6KAU8 | ATG16L2 | 18 | 100.000 | 3.292 | 100.000 | 156.16 |
| Q9CQY1 | ATG12 | 5 | 44.585 | 15.343 | 2.043 | 68.65 |
| Q9CPX6 | ATG3 | 16 | 43.193 | 100.000 | 0.010 | 351.60 |
| P60521 | GABARAPL2 | 6 | 38.693 | 100.000 | 0.010 | 200.07 |
| Q80VP0 | TECPR1 | 37 | 26.344 | 0.975 | 40.957 | 519.22 |
| Q3TAB9 | ATG7 | 22 | 22.541 | 100.000 | 0.152 | 122.13 |
| Q8BJL1 | FBXO30 | 2 | 19.013 | 42.700 | 0.320 | 2.26 |
| Q99J83 | ATG5 | 18 | 18.431 | 0.640 | 73.999 | 690.24 |
| P62192 | PSMC1 | 3 | 11.558 | 1.838 | 6.287 | 14.06 |
| Q9DAX9 | APPBP2 | 3 | 7.636 | 0.371 | 20.567 | 8.94 |
| Q68FD5 | CLTC | 50 | 7.517 | 11.637 | 0.665 | 310.55 |
| Q9QZZ6 | DPT | 3 | 5.980 | 0.294 | 12.435 | 19.01 |
| E9Q9A9 | OAS2 | 20 | 5.926 | 0.530 | 100.000 | 55.25 |
| Q3UIZ0 | GAK | 12 | 5.614 | 25.220 | 0.405 | 27.14 |
| Q03350 | THBS2 | 5 | 4.939 | 0.530 | 67.590 | 20.45 |
| Q3TJ69 | SERPINB9B | 6 | 4.919 | 1.652 | 1.645 | 12.80 |
| P39053 | DNM1 | 2 | 4.848 | 3.489 | 3.567 | 30.78 |
| B1AQR8 | LGALS9 | 4 | 4.761 | 3.392 | 1.187 | 6.31 |
| E9QP46 | SYNE2 | 2 | 4.325 | 100.000 | 0.010 | 4.39 |
| B2RXT5 | GPI1 | 3 | 4.047 | 0.835 | 5.621 | 4.25 |
| B1AWE0 | CLTA | 2 | 4.040 | 8.772 | 0.461 | 4.04 |
| P61804 | DAD1 | 3 | 3.952 | 0.922 | 0.788 | 18.58 |
| F8VPL2 | PIK3C2A | 10 | 3.836 | 11.818 | 0.300 | 19.49 |
| Q3TCY0 | RAB33B | 4 | 3.736 | 5.842 | 0.628 | 10.01 |
| P10649 | GSTM1 | 2 | 3.368 | 1.214 | 2.775 | 3.97 |
| P62814 | ATP6V1B2 | 3 | 3.293 | 1.494 | 1.875 | 9.26 |
| O35646 | CAPN6 | 8 | 2.952 | 0.909 | 3.153 | 22.75 |
| A2RSJ4 | UHRF1BP1L | 2 | 2.906 | 91.980 | 0.032 | 4.14 |
| B9EJ77 | TANC1 | 4 | 2.902 | 2.167 | 1.398 | 9.96 |
| P63154 | CRNKL1 | 2 | 2.887 | 11.270 | 0.083 | 5.09 |
| Q9CZK0 | SNX9 | 7 | 2.599 | 3.712 | 0.687 | 20.79 |
| P41731 | CD63 | 2 | 2.578 | 2.932 | 1.000 | 1.81 |
| Q3U1H7 | SNX18 | 2 | 2.555 | 3.208 | 0.796 | 1.62 |
| Q5NCB5 | IRGM1 | 3 | 2.553 | 0.761 | 4.725 | 11.45 |
| Q3TRC8 | ARRB2 | 2 | 2.549 | 2.637 | 0.967 | 1.88 |
| M0QW74 | MTMR3 | 2 | 2.521 | 100.000 | 0.013 | 6.81 |
| Q3T9X3 | DNM2 | 8 | 2.500 | 2.710 | 0.917 | 48.27 |
| B2RRF0 | PTPRK | 3 | 2.427 | 0.741 | 44.851 | 7.89 |
| Q60575 | KIF1B | 3 | 2.381 | 1.181 | 3.083 | 22.94 |
| Q9DC42 | CD97 | 5 | 2.372 | 2.666 | 0.794 | 11.63 |
| Q921W0 | CHMP1A | 2 | 2.336 | 0.817 | 2.859 | 6.79 |
| Q8BIF7 | GRWD1 | 2 | 2.316 | 17.317 | 0.086 | 3.90 |

| | | | | | | |
|---|---|---|---|---|---|---|
| Q80UP1 | HIP1R | 3 | 2.311 | 81.993 | 0.028 | 3.35 |
| Q9D154 | SERPINB1A | 4 | 2.262 | 0.618 | 7.558 | 5.00 |
| Q69ZA7 | MKIAA1769 | 2 | 2.248 | 1.473 | 1.526 | 7.98 |
| O08797 | SERPINB9 | 4 | 2.202 | 0.696 | 2.814 | 20.40 |
| Q9QYR6 | MAP1A | 3 | 2.157 | 1.281 | 1.341 | 10.87 |
| Q61749 | EIF2B4 | 3 | 2.130 | 2.184 | 1.115 | 14.16 |
| Q9CQV8 | YWHAB | 5 | 2.114 | 1.804 | 1.142 | 113.95 |
| E9Q555 | RNF213 | 113 | 2.110 | 1.379 | 1.568 | 527.03 |
| Q9WTQ5 | AKAP12 | 13 | 2.086 | 2.930 | 0.606 | 28.85 |
| B2RQS1 | STRN3 | 3 | 2.074 | 1.089 | 1.904 | 4.44 |
| Q5MJ56 | MYO7A | 2 | 2.062 | 2.789 | 0.631 | 4.27 |
| Q9DB90 | SMG9 | 2 | 2.044 | 3.343 | 0.586 | 4.39 |
| Q8K234 | OASL2 | 7 | 2.034 | 0.499 | 3.447 | 31.04 |
| Q3TW11 | STAT1 | 25 | 2.030 | 0.962 | 2.197 | 138.03 |
| P11928 | OAS1A | 4 | 2.018 | 0.638 | 2.664 | 13.89 |
| Q9ER88 | DAP3 | 2 | 2.005 | 1.533 | 1.484 | 3.36 |

**Supplemental Table 5:** GFP-ATG5 interactome in the ATG3 null background. Putative interactors represented by 2 or more unique peptides ranked in order of: (i) wild type GFP-ATG5:GFP interactors (>2-fold enrichment); (ii) wild-type:mutant GFP-ATG5 interactors (>2-fold enrichment); (iii) score. Autophagy proteins are highlighted green; membrane trafficking proteins are highlighted yellow.

| *Increased* | | | | *Decreased* | | | |
|---|---|---|---|---|---|---|---|
| **Accession** | **Description** | **Mean WT : GFP** | ***P*-value** | **Accession** | **Description** | **Mean WT : GFP** | ***P*-value** |
| P52800 | EFNB2 | 2.730 | 0.035033748 | P70202 | LXN | 0.564 | 0.030098094 |
| Q9JKF6 | PVRL1 | 2.316 | 0.010530861 | Q9R0P5 | DSTN | 0.595 | 0.017489458 |
| A2A8L5 | PTPRF PE | 2.205 | 0.022140881 | A0A067XG53 | CASK | 0.597 | 0.015246012 |
| Q9QXX0 | JAG1 | 2.044 | 0.002498732 | Q05DU8 | RRM1 | 0.605 | 0.028377966 |
| Q3UXH8 | HDAC2 | 1.765 | 0.018127927 | Z4YL78 | CKAP5 | 0.616 | 0.018539603 |
| Q61876 | Uncharacterized | 1.649 | 0.034883794 | Q9QYF9 | NDRG3 | 0.634 | 0.02342693 |
| Q3TDB6 | SLC31A1 | 1.533 | 0.034144225 | Q6QHF0 | TNFRSF1A | 0.640 | 0.022715216 |
| Q9CSH0 | HNRNPLL | 1.445 | 0.005323671 | Q91VE0 | SLC27A4 | 0.641 | 0.032681628 |
| Q3TDN0 | DISP1 | 1.434 | 0.018178448 | F8VQD7 | PTPRG | 0.642 | 0.017172075 |
| Q3TJG5 | SLC12A4 | 1.428 | 0.002214814 | Q64337 | SQSTM1 | 0.662 | 0.045255171 |
| Q6PB66 | LRPPRC | 1.421 | 0.022707129 | P27046 | MAN2A1 | 0.664 | 0.044194909 |
| P62814 | ATP6V1B2 | 1.420 | 0.0041942 | Q3UDS4 | SQRDL | 0.699 | 0.016460184 |
| Q3U449 | BPNT1 | 1.379 | 0.032157651 | Q9DBT5 | AMPD2 | 0.731 | 0.03829524 |
| Q8BFY6 | PEF1 | 1.323 | 0.012362173 | P70302 | STIM1 | 0.755 | 0.042594089 |
| A0A0A0U6W1 | Envelope protein | 1.317 | 0.044056924 | P47738 | ALDH2 | 0.758 | 0.041949934 |
| | | | | O09172 | GCLM | 0.762 | 0.044379611 |

**Supplemental Table 7:** Surface interactome in WT GFP-ATG5 vs. GFP in the fed state. These represent the proteins whose expression is increased (green shading) or decreased (orange shading) >1.3 fold with $p < 0.05$.

| *Increased* | | | | *Decreased* | | | |
|---|---|---|---|---|---|---|---|
| **Accession** | **Description** | **Mean WT : K130R** | ***P*-value** | **Accession** | **Descripti on** | **Mean WT : K130R** | ***P*-value** |
| E9PVD3 | DCHS1 | 2.334 | 0.0016836 | E9PV48 | IFIT3B | 0.013 | 0.00094788 |
| P98156 | VLDLR | 2.063 | 0.04272076 | F6VQ81 | TPD52L2 | 0.551 | 0.03955201 |
| P70207 | PLXNA2 | 1.917 | 0.03147214 | Q91VE0 | SLC27A4 | 0.600 | 0.03470227 |
| Q8BP67 | RPL24 | 1.759 | 0.01573821 | Q62087 | PON3 | 0.607 | 0.00355953 |
| O35607 | BMPR2 | 1.739 | 0.02682914 | P27046 | MAN2A1 | 0.620 | 0.01859173 |
| A2ATK9 | FAM171A1 | 1.718 | 0.02862772 | G3UXZ5 | PSME1 | 0.645 | 0.01229071 |
| P62754 | RPS6 | 1.708 | 0.00477564 | Q9CQ43 | DUT | 0.648 | 0.04658237 |
| Q91VD8 | PCDHB17 | 1.680 | 8.3049E-05 | Q01730 | RSU1 | 0.653 | 0.03962807 |
| E9QM38 | SLC12A2 | 1.663 | 0.02232769 | P45952 | ACADM | 0.662 | 0.01518668 |
| Q9D6G1 | HNRNPAB | 1.621 | 0.00407523 | P70302 | STIM1 | 0.672 | 0.0484652 |
| O89051 | ITM2B | 1.620 | 0.04344145 | O78207 | H2-D1 | 0.674 | 0.01256854 |
| Q99KP6 | PRPF19 | 1.619 | 0.0285495 | P26043 | RDX | 0.685 | 0.03325535 |
| Q7TT36 | ADGRA3 | 1.574 | 0.02674282 | P70303 | CTPS2 | 0.689 | 0.04017958 |
| Q60737 | CSNK2A1 | 1.566 | 0.00374756 | Q8BJY1 | PSMD5 | 0.702 | 0.03550639 |
| Q8VIK5 | PEAR1 | 1.552 | 0.02106129 | P97429 | ANXA4 | 0.704 | 0.01723249 |
| A2A699 | FAM171A2 | 1.538 | 0.04764387 | O88587 | COMT | 0.705 | 0.03218573 |
| E9PUQ9 | PIEZO1 | 1.450 | 0.02592485 | Q9D071 | MMS19 | 0.7076 | 0.01865301 |
| Q3U561 | RPL10A | 1.494 | 0.0203584 | A0A0R4J0H7 | NCAPD2 | 0.709 | 0.01514302 |
| Q9D074 | MGRN1 | 1.485 | 0.01024261 | E9Q855 | SCAMP3 | 0.712 | 0.0137413 |
| Q8C0I1 | AGPS | 1.458 | 0.04866581 | F8VQC1 | SRP72 | 0.727 | 0.04605419 |
| Q3TF81 | RPP30 | 1.437 | 0.02715007 | P31938 | MAP2K1 | 0.733 | 0.00360613 |
| Q61090 | FZD7 | 1.396 | 0.01208308 | P26039 | TLN1 | 0.734 | 0.04548505 |
| G3UYZ1 | IGSF8 | 1.361 | 0.00133379 | O35685 | NUDC | 0.747 | 0.0295591 |
| Q9WV91 | PTGFRN | 1.342 | 0.01585211 | Q99JW7 | CDK1 | 0.754 | 0.02838882 |
| P97792 | CXADR | 1.330 | 0.03203783 | | | | |
| Q8BKG3 | PTK7 | 1.330 | 0.01252472 | | | | |
| E9PXY1 | CUL4B | 1.322 | 0.01816524 | | | | |
| P82347 | SGCD | 1.317 | 0.04105339 | | | | |

**Supplemental Table 8:** Surface interactome in WT GFP-ATG5 vs. K130R GFP in the fed state. These represent the proteins whose expression is increased (green shading) or decreased (orange shading) >1.3 fold with $p < 0.05$.

***Increased***

| Accession | Description | Mean K130R : GFP | *P*-value |
|---|---|---|---|
| Q9JHJ8 | ICOSLG | 3.611 | 0.031519 |
| B2RWZ0 | ABI3BP | 2.341 | 0.020331 |
| Q9CQ62 | DECR1 | 1.879 | 0.0399712 |
| Q99J39 | MLYCD | 1.813 | 0.0099304 |
| Q3UAG2 | PGD | 1.634 | 0.0203635 |
| P08030 | APRT | 1.566 | 0.0261245 |
| Q9QXB9 | DRG2 | 1.560 | 0.0402071 |
| Q62186 | SSR4 | 1.444 | 0.0066468 |
| Q8BGX2 | C19ORF52 | 1.435 | 0.023599 |
| Q8C5P5 | NT5DC1 | 1.4172 | 0.015404 |
| Q9CQ43 | DUT | 1.338 | 0.0447269 |
| P35700 | PRDX1 | 1.330 | 0.028688 |
| Q3U8R9 | TXNL1 | 1.328 | 0.0287701 |
| A0A0U1RNT6 | AUH | 1.305 | 0.048153 |

***Decreased***

| Accession | Description | Mean K130R : GFP | *P*-value |
|---|---|---|---|
| P57787 | SLC16A3 | 0.439 | 0.0299909 |
| P02469 | LAMB1 | 0.516 | 0.0196555 |
| Q3TJH1 | GNAI3 | 0.597 | 0.0474658 |
| Q6PDG0 | NUP205 | 0.599 | 0.0424557 |
| P40240 | CD9 | 0.599 | 0.0345749 |
| A0A0R4J097 | TGFBR3 | 0.605 | 0.0305692 |
| Q8R373 | CLMP | 0.649 | 0.0066052 |
| Q8BLU0 | FLRT2 | 0.656 | 0.0163612 |
| F8VQJ3 | LAMC1 | 0.668 | 0.0400735 |
| Q8BUM1 | TARDBP | 0.682 | 0.0066638 |
| F8VQD7 | PTPRG | 0.689 | 0.0383892 |
| Q9QYF9 | NDRG3 | 0.699 | 0.0481767 |
| Q9DBV4 | MXRA8 | 0.701 | 0.0287062 |
| A2ATK9 | FAM171A1 | 0.702 | 0.038105 |
| Q3TCZ2 | SLC29A1 | 0.708 | 0.0211726 |
| P70424 | ERBB2 | 0.711 | 0.0337593 |
| O35188 | CX3CL1 | 0.721 | 0.0369577 |
| Q61090 | FZD7 | 0.725 | 0.0225818 |
| A0A0R4J0A9 | LRP6 | 0.751 | 0.0082895 |
| Q8BKG3 | PTK7 | 0.755 | 0.0053365 |
| P97351 | RPS3A | 0.761 | 0.0309467 |

**Supplemental Table 9:** Surface interactome in K130R GFP vs GFP in the fed state. These represent the proteins whose expression is increased (green shading) or decreased (orange shading) >1.3 fold with $p < 0.05$.

#### WT GFP-ATG5 vs. GFP

| *Increased* | | | | *Decreased* | | | |
|---|---|---|---|---|---|---|---|
| **Accession** | **Description** | **Mean** | ***P*-value** | **Accession** | **Description** | **Mean** | ***P*-value** |
| Q3V3R1 | MTHFD1L | 1.506 | 0.037687061 | E9PX70 | COL12A1 | 0.401 | 0.016454101 |
| G5E843 | ROBO1 | 1.483 | 0.025731268 | Q8R2Q8 | BST2 | 0.469 | 0.02475123 |
| A2A813 | PARK7 | 1.417 | 0.01063417 | Q8CE18 | Uncharacterised | 0.564 | 0.013425909 |
| Q8R464 | CADM4 | 1.370 | 0.049862472 | Q3TVX7 | SYPL | 0.62 | 0.031614541 |
| Q9R0B9 | PLOD2 | 1.340 | 0.047553399 | Q9CQW9 | IFITM3 | 0.63 | 0.046282885 |
| F7BWT7 | TSPAN15 | 1.307 | 0.003948324 | O78207 | H2-D1 | 0.632 | 0.041047422 |
| Q3THB3 | HNRNPM | 1.304 | 0.018914629 | Q3TWF3 | APP | 0.65 | 0.043893734 |
| | | | | G3UXZ5 | PSME1 | 0.663 | 0.042184701 |
| | | | | P11438 | LAMP1 | 0.665 | 0.032615735 |
| | | | | O35316 | SLC6A6 | 0.674 | 0.037689164 |
| | | | | Q3TDG9 | STX12 | 0.758 | 0.01266362 |

#### WT GFP-ATG5 vs. K130R GFP-ATG5

*Decreased*

| Accession | Description | Mean | *P*-value | Accession | Description | Mean | *P*-value |
|---|---|---|---|---|---|---|---|
| P11438 | LAMP1 | 0.389 | 0.00064169 | P24369 | PPIB | 0.690 | 0.043396898 |
| Q8R2Q8 | BST2 | 0.489 | 0.018887447 | Q9R1R9 | RDH11 | 0.699 | 0.014634982 |
| Q8BLN5 | LSS | 0.539 | 0.015667318 | Q9D0F3 | LMAN1 | 0.701 | 0.022609471 |
| Q3TWF3 | APP | 0.632 | 0.000819975 | Q6ZQI3 | MLEC | 0.709 | 0.04219749 |
| Q61398 | PCOLCE | 0.637 | 0.030960639 | Q80UU9 | PGRMC2 | 0.723 | 0.039208203 |
| E9QMJ5 | ADGRE5 | 0.645 | 0.04195319 | Q05DV1 | POR | 0.733 | 0.001021338 |
| Q6PB52 | LRPAP1 | 0.646 | 0.036486923 | Q61833 | RPN2 | 0.744 | 0.00793402 |
| Q91VE0 | SLC27A4 | 0.648 | 0.011679863 | Q08879 | FBLN1 | 0.746 | 0.044594835 |
| P27773 | PDIA3 | 0.658 | 0.049089537 | O70503 | HSD17B12 | 0.762 | 0.005339892 |
| Q9QXT0 | CNPY2 | 0.660 | 0.038651952 | O08807 | PRDX4 | 0.767 | 0.035518398 |
| Q3TDN2 | FAF2 | 0.669 | 0.011108781 | Q5NCU4 | SPARC | 0.767 | 0.026942681 |

#### K130R GFP-ATG5 vs. GFP

| *Increased* | | | | *Decreased* | | | |
|---|---|---|---|---|---|---|---|
| **Accession** | **Description** | **Mean** | ***P*-value** | **Accession** | **Description** | **Mean** | ***P*-value** |
| P11438 | LAMP1 | 1.920 | 0.025481169 | P11438 | MOB1B | 0.660 | 0.006370983 |
| Q9EP69 | SAC1 | 1.783 | 0.0451057 | Q9EP69 | PPIC | 0.674 | 0.049442382 |
| G3UVV4 | HK1 | 1.685 | 0.001635713 | G3UVV4 | RPL7 | 0.716 | 0.024964515 |
| Q3V3R1 | MTHFD1L | 1.666 | 0.012400546 | Q3V3R1 | RPL28 | 0.718 | 0.012843299 |
| A1L2Z3 | EMC1 | 1.526 | 0.044983442 | A1L2Z3 | CAMK2D | 0.746 | 0.014498686 |
| P46978 | STT3A | 1.471 | 0.006492825 | P46978 | ACACA | 0.750 | 0.027215708 |
| Q8BVQ5 | PPME1 | 1.442 | 0.021765395 | Q8BVQ5 | ERBB2 | 0.758 | 0.005412206 |
| Q8BKE6 | CYP20A1 | 1.438 | 0.023491593 | Q8BKE6 | IL1RAP | 0.758 | 0.014312666 |
| P58242 | SMPDL3B | 1.400 | 0.023246228 | P58242 | RUVBL1 | 0.763 | 0.049164754 |

| | | | |
|---|---|---|---|
| Q99KI0 | ACO2 | 1.337 | 0.020549282 |
| Q3U7R1 | ESYT1 | 1.307 | 0.025087592 |

**Supplemental Table 11:** Surface interactome in starved MEF rescue backgrounds. Proteins increased (top; green shading) or decreased (bottom; orange shading) >1.3 fold with $p < 0.05$ are shown.

#### WT GFP-ATG5 vs. GFP

| *Increased* | | | | *Decreased* | | | |
|---|---|---|---|---|---|---|---|
| **Accession** | **Description** | **Starved/Fed mean** | ***P*-value** | **Accession** | **Description** | **Starved/Fed mean** | ***P*-value** |
| Q9R0P5 | DSTN | 1.775 | 0.020129119 | Q3TVX7 | SYPL | 0.499202051 | 0.021841626 |
| A2A813 | PARK7 | 1.617 | 0.044781611 | P11438 | LAMP1 | 0.579 | 0.004855501 |
| Q9R0B9 | PLOD2 | 1.473 | 0.025822975 | P04925 | PRNP | 0.664 | 0.046346883 |
| O09172 | GCLM | 1.323 | 0.027309868 | | | | |

#### WT GFP-ATG5 vs. K130R GFP-ATG5

| *Increased* | | | | *Decreased* | | | |
|---|---|---|---|---|---|---|---|
| **Accession** | **Description** | **Starved/Fed mean** | ***P*-value** | **Accession** | **Description** | **Starved/Fed mean** | ***P*-value** |
| Q9WVG6 | CARM1 | 1.707 | 0.027837244 | Q9EPR5 | SORCS2 | 0.377 | 0.033646884 |
| Q9CQ65 | MTAP | 1.590 | 0.038584358 | P53994 | RAB2A | 0.641 | 0.01405476 |
| E9PVA8 | GCN1 | 1.589 | 0.040604744 | Q05DV1 | POR | 0.695 | 0.023477359 |
| Q8CG48 | SMC2 | 1.537 | 0.036123316 | | | | |
| P26039 | TLN1 | 1.448 | 0.030744401 | | | | |
| Q9JKF1 | ICGAP1 | 1.379 | 0.037147102 | | | | |

#### K130R GFP-ATG5 vs. GFP

| *Increased* | | | | *Increased* | | | |
|---|---|---|---|---|---|---|---|
| **Accession** | **Description** | **Starved/Fed mean** | ***P*-value** | **Accession** | **Description** | **Starved/Fed mean** | ***P*-value** |
| Q8BKE6 | CPY20A1 | 1.498 | 0.045389404 | P19096 | FASN | 0.687 | 0.039200995 |

**Supplemental Table 12:** Surface interactome changes upon starvation (ratio starved:fed state) in the different MEF rescue backgrounds. Proteins increased (top; green shading) or decreased (bottom; orange shading) >1.3 fold with p< 0.05 are shown.
